# scDenorm: a denormalisation tool for integrating single-cell transcriptomics data

**DOI:** 10.1101/2025.05.10.653289

**Authors:** Yin Huang, Anna Vathrakokili Pournara, Ying Ao, Lirong Yang, Hui Zhang, Yongjian Zhang, Sheng Liu, Alvis Brazma, Irene Papatheodorou, Xinlu Yang, Ming Shi, Zhichao Miao

## Abstract

Integrating single-cell omics data at an atlas scale enhances our understanding of cell types and disease mechanisms. However, the integration of data processed by different normalisation methods can lead to biases, such as unexpected batch effects and gene expression distortion, leading to misinterpretations in downstream analysis. To address these challenges, we present scDenorm, an algorithm that reverts normalised single-cell omics data to raw counts, preserving the integrity of the original measurements and ensuring consistent data processing during integration. We evaluated scDenorm’s performance on large-scale datasets and benchmarked its impact on data integration and downstream analysis across three datasets.

## Background

Single-cell RNA sequencing (scRNA-seq) is a powerful high-throughput technology for measuring gene expression in individual cells. Integration of atlas-level single-cell transcriptomics data has exerted a great potential for understanding how cells orchestrate in the human body, as well as complex molecular mechanisms in various diseases^1,2^. With the progress of the Human Cell Atlas (HCA)^3^, an increasing number of reference atlases are available for comparison and integration^4–6^. Numerous integration methods have been developed and several studies have been performed to benchmark their performance and explore their limitations^7–9^. In order to achieve effective large-scale data integration, it is crucial to take into account the assumptions of data distribution and noise levels. For instance, scVI^10^ and scANVI^11^ integration methods model single-cell data using a Negative Binomial distribution (also known as Gamma-Poisson distribution), and thus both require raw counts as input. Even though some other integration methods, *e*.*g*., Seurat integration methods^12^ (RPCA, CCA), Harmony^13^, Liger^14^ etc, do not directly rely on raw counts, they inherently make assumptions about the data distribution. As a result, for most existing integration methods, it is key to ensure the consistency of the input datasets.

To address technical variations (*e*.*g*., sequencing depth) and biases inherent in scRNA-seq, scaling and transformation methods are often employed to ensure comparability across cells^15,16^. Normally, scaling is used to account for sequencing depth, while transformation is used to stabilise variance of the data. The differences between variance-stabilising transformations have been benchmarked by Constantin Ahlmann-Eltze and Wolfgang Huber^17^, demonstrating the effectiveness of the delta method for comparing cells with varying gene expression levels. In a delta normalisation, raw counts are scaled by total counts and target sum, followed by log-transformation with an added pseudo count (see **Methods**). It has been adopted in well-established analysis workflows (e.g., Seurat^18^ and SCANPY^19^), assuming that droplet-based scRNA-seq data follows a negative binomial distribution^20–23^. In some large-scale data resources, such as the UCSC Cell Browser^24^, delta method normalised matrices are deposited instead of the raw counts to facilitate reproducibility of analysis results. Thus, many datasets are available only as processed matrices rather than raw counts, hindering atlas-level data integration.

The best way to guarantee consistent data processing in large-scale data integration is to use raw counts as input. Integrating processed data with raw counts can introduce unnecessary biases, due to re-normalising of data that has already been normalised. Some downstream analysis steps^25,26^ (such as selecting highly variable genes, differential gene expression analysis) also assume raw counts as input. When raw counts are not available, researchers often seek to obtain the raw sequencing data and re-analyse them, including secondary analysis of reads mapping, demultiplexing, and quantification analysis^27^ to obtain the raw count matrix. However, a count matrix from the re-analysis may deviate from the original published analysis in terms of reference genome and cell barcodes. Thus, the cell type annotation or other metadata reported in the raw publication cannot be used, rendering difficulties in reproducing the analysis results. Besides, this secondary analysis can be both computationally expensive and time-consuming. Therefore, reliable conversion of normalised matrices back to raw counts can benefit large-scale data integration tasks as well as wider use of publicly deposited data. Yet, there is no tool available to meet this urgent need.

In this study, we propose scDenorm, an algorithm that converts delta method normalised gene expression data back to the raw counts. It effectively explores key implicit features of the data distribution in scRNA-seq and recovers raw count matrices. Based on benchmarking across large-scale datasets, as well as application studies of downstream analysis, we demonstrate the capability, accuracy, scalability and efficiency of this method. Moreover, scDenorm can deal with different normalisation parameters, thereby facilitating data integration, consistent downstream analyses and the construction of atlases.

## Results

### Inconsistent data normalisation may generate biases in data integration

Using the 10x 3k peripheral blood mononuclear cells (PBMCs) data, which is the example data used in the well-established SCANPY^19^ and Seurat^18^ single-cell tutorials, as an example, we investigated the impact of normalisation parameters in the delta method. These parameters are the scaling factor, logarithmic transformation base, and pseudo counts. Matrices normalised by different parameters go through the same downstream analysis of highly variable gene selection, dimensionality reduction (e.g., Principal Component Analysis), clustering and visualisation. The Uniform Manifold Approximation and Projection (UMAP) plot shows deviations between datasets processed with different normalisation parameters, e.g., the deviations between B cells in L=10^3^ and the same B cells in other normalisations, indicating the potential bias introduced by inconsistent data normalisation (**Fig. 1a,b, Supplementary Fig. 1a**). Furthermore, such a data normalisation effect cannot be removed through data integration by Harmony^13^, scanorama^28^, or BBKNN^29^ (**Fig. 1, Supplementary Fig. 1**), e.g., the B cell populations in **Fig. 1c** and **1d** cluster separately. Therefore, we suggest converting the normalised matrices back to raw counts for a consistent data integration and downstream analysis.

**Fig. 1.**
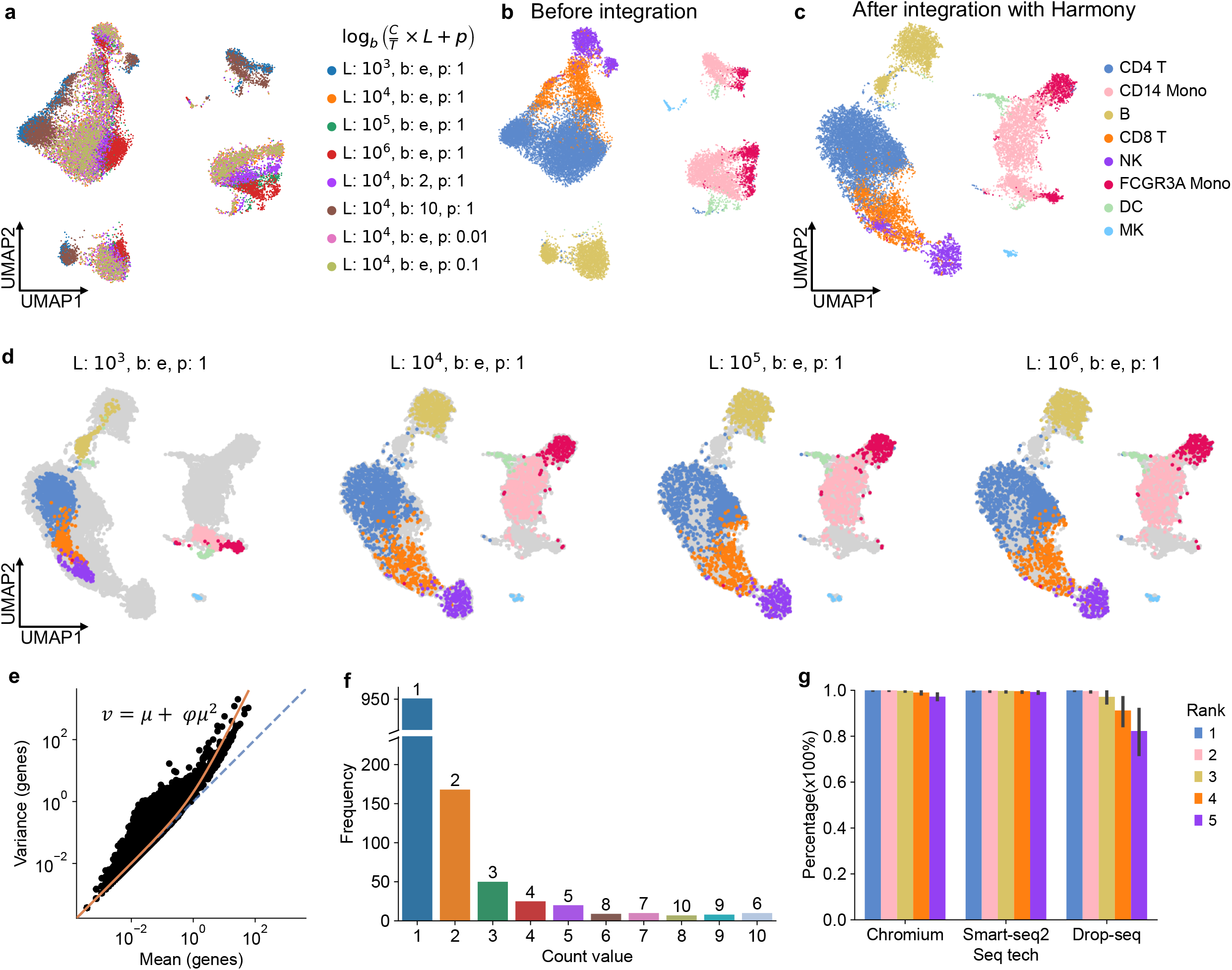
The data distribution of droplet-based single-cell data **a**, UMAP plot of PBMC 3k datasets, without data integration, normalised with different delta normalisation parameters, including target sum (L), logarithmic base (b) and pseudo count (p). The plot is coloured by different parameter sets. **b**, The same UMAP plot as panel (**a**) coloured according to cell type annotation. **c**, UMAP plot after Harmony integration of data normalised by different parameters, coloured by cell types. **d**, The UMAP plots are the same as panel (**c**) (after data integration by Harmony), displaying four different normalisation parameter sets. **e**, Scatter plot demonstrating the mean (x-axis) against the variance (y-axis) for each gene in the count matrix of the PBMC 3k datasets. Each dot shows the mean and variance value of a gene. The diagonal line is shown in blue. The orange curve is the fitted curve of negative binomial distribution with variance *v*, mean μ and dispersion φ. **f**, Histogram depicting the frequencies of count values and their ranks in a single cell, showing the ‘count-rank’ distribution in a cell selected from the count matrix of the PBMC 3k datasets. **g**, The percentage of cells that follow the ‘count-rank’ distribution (the value of count equal to its rank from 1 to 5) in three scRNA-seq technologies (Chromium, Smart-seq2, and Drop-seq).

### The denormalisation process in scDenorm

We term the recovery of normalised data to raw counts as “denormalisation”. Denormalising delta method normalised data requires the determination of three parameters, scaling factors, the logarithmic transformation (log-transformation) base, and the pseudo count. The first step for denormalisation is to determine if log-transformed has been applied for the whole expression matrix. It is well established that droplet-based scRNA-seq data follows a negative binomial distribution^20–23^, where the variance exceeds the mean (**Fig. 1e**). Thus, the variance versus mean distribution effectively indicates whether the data has been log-transformed or not (**Supplementary Fig. 2a**). The second key step in denormalisation is to determine the scaling factor for each cell. This needs to exploit the implicit data distribution feature of scRNA-seq. Droplet-based scRNA-seq mainly probes the highly expressed genes, rendering a high dropout rate. In a sparse matrix where zeros have been removed, the frequency of counts can be ranked, with the most frequent count number being one, followed by two, and so on (**Fig. 1f, Supplementary Fig. 2b,c**). Using such a ‘count-rank’ distribution, scaling factors for cells can be measured by establishing the relationship between the top two most frequent numbers in the normalised data and numbers one and two. After exploring 105 datasets from the Brain Cell Atlas^30^, we found that over 99% of the cells follow this ‘count-rank’ distribution for the top three most frequent count numbers (one, two, and three), while >95% cells in Chromium and > 80% cells in Drop-seq follow the distribution for the top five count numbers. Notably, >99% cells in Smart-seq2 data follow this distribution for the top ten numbers (**Fig. 1g, Supplementary Fig. 2d**). Following this ‘count-rank’ distribution, the top most frequent count numbers can be used to determine the three parameters in delta method normalisation (**Supplementary Fig. 3a**).

The denormalisation procedure in scDenorm involves two steps: de-transformation and unscaling (**Supplementary Fig. 3b**). In the de-transformation step, a subset matrix (100 cells) is used to determine the same log-transformation base and pseudo count among cells since these two parameters keep the same for the whole expression matrix. Using a subset of data effectively accelerates the calculation. First, empirical values (*e*.*g*., 2, e Euler’s number, 10 for the log-transformation base, 0.01, 0.1, 1 for pseudo count), which are used in standard analysis workflows, are tried. If not successful, these two parameters can be determined by solving equations between the top two most frequent numbers (**Supplementary Fig. 3c**). In the unscaling step, each cell has a different scaling factor, which is a ratio between the total counts of the cell and the target sum (e.g., 10,000). To measure the scaling factor of a cell, we implement two methods: 1) a regression-based method (see **Methods**, equation (4) in **Supplementary Fig. 3d**) and 2) solving equations between the top two most frequent numbers (see **Methods**, equation (5) in **Supplementary Fig. 3d**), while the latter method offers the advantages of fast speed and good robustness (**Supplementary Fig. 4a,b**). As the expression matrix is processed from raw counts, which consists of integers only, a successful denormalisation should result in a small mean square error between denormalised values and their nearest integers (see **Methods**).

To elaborate the denormalisation process, we used an example dataset^31^ of single-nucleus RNA sequencing (snRNA-seq) data of autism spectrum disorder, with both the normalised data and raw count matrix available in the database from https://autism.cells.ucsc.edu. According to the respective publication^31^, the data was normalised with the delta method. The relationship between the top ten most frequent gene expression values and their respective frequencies in three cells in the processed data (**Fig. 2a**), suggests a logarithmic distribution, while the less frequent values after them do not follow such a distribution due to dropouts. Should any of the cells in the dataset follow the same distribution. And the mean versus variance distribution confirms a logarithmic conversion (**Fig. 2b**). The log-transformation base and the pseudo count are determined as 2 and 1 respectively, by solving equation (3) in **Supplementary Fig. 3c**. These parameters show a good fit according to the top two most frequent values (**Fig. 2c**). The normalised matrix is de-transformed by taking the exponential of the log-transformation base and subtracting the pseudo count, resulting in a ‘scaled matrix’. In the scaled matrix, the top five most frequent values show a linear ‘count-rank’ distribution in each cell (**Fig. 2d**). The slope of the line is the reciprocal of the scaling factor. This linear distribution indicates the success of de-transformation. Additionally, the mean versus variance distribution (**Fig. 2e**) confirms this success. The summed expression values for most cells are approximately 10,000, indicating that the target sum is 10,000. Some genes may have been removed after normalisation, leading to a reduction in the summed expression values (**Fig. 2f**). In the unscaling step, scaling factors are determined by solving equation (5) in **Supplementary Fig. 3d**. Each cell is multiplied by its scaling factor resulting in a “denormalised matrix”, which is supposed to be similar to the raw count matrix of integers. As in a sparse matrix, the top two most frequent numbers should be one and two (**Fig. 2g**). The mean versus variance distribution of the denormalised matrix conforms to a negative binomial distribution (**Fig. 2h**), which is expected for the raw counts of droplet-based scRNA-seq. Comparing the denormalised matrix with the raw count matrix, the maximum error for each value was less than 0.001 (**Fig. 2i**), which may result from the digital float calculation. After taking round values, the denormalised matrix is identical to the raw count matrix, suggesting a successful denormalisation.

**Fig. 2.**
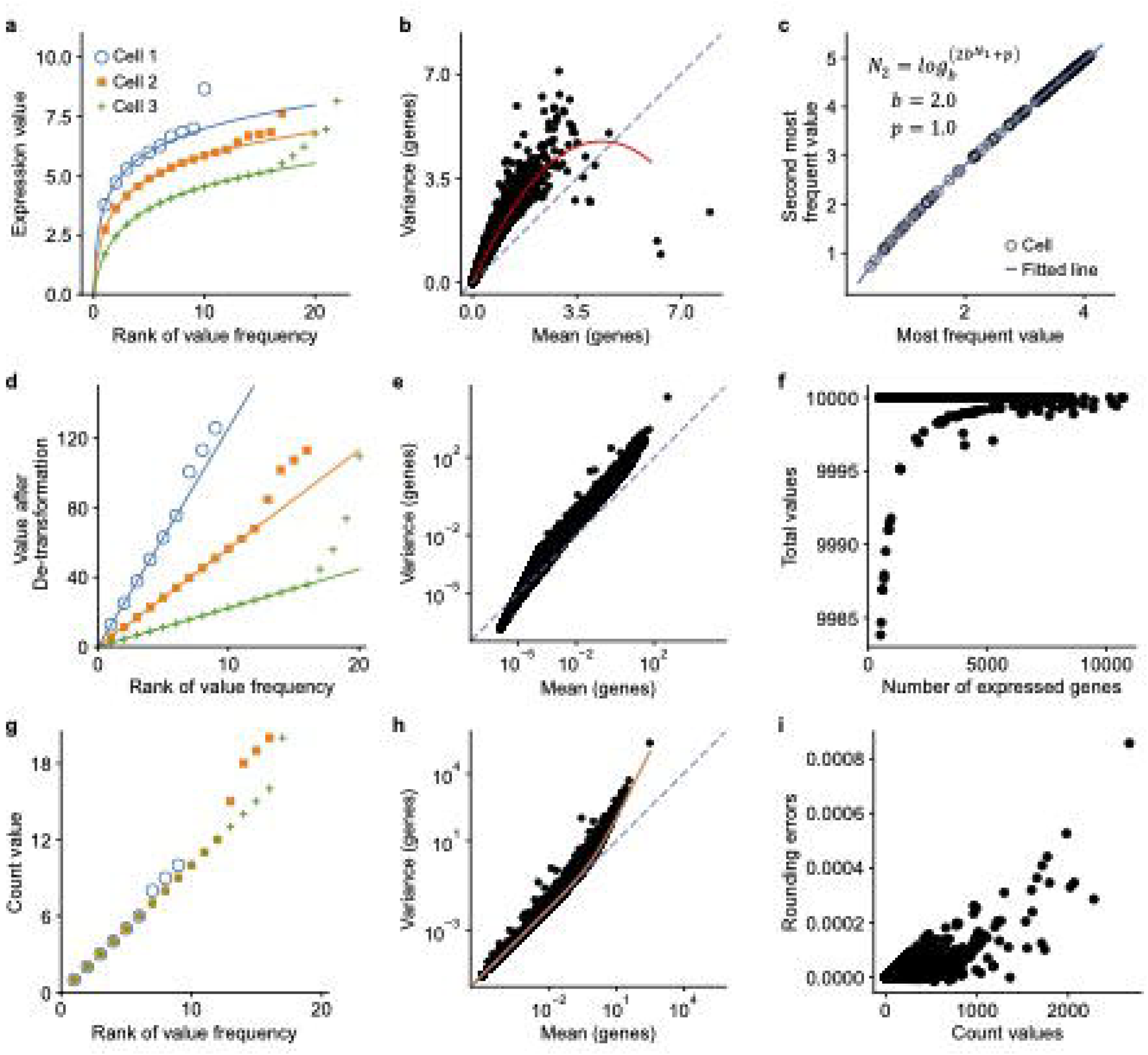
Evaluation of scDenorm on normalised scRNA-seq data with known raw counts **a**, The scatter plot shows the distribution between expression values and their ranks of frequencies in three example cells. Each dot is an expression value and its rank of frequency in the cell, different cells are shown in different shape and colour. **b**, the scatter plot shows the distribution between the log-transformed mean expression (x-axis) and the log-transformed variance (y-axis) for each gene in the gene expression matrix from Velmeshev et al dataset. The diagonal line (x=y) is shown in blue. **c**, The scatter plot shows the distribution between the most and second most frequent values in different cells, displaying each cell as a dot. The blue curve shows the fitted equation derived from the equation (4) in **Methods**, with base value (b) equal to 2 and pseudo count (p) equal to 1. N_1_ and N_2_ are the most and second most frequent values in cells. **d**, The scatter plot shows the distribution between expression values and their ranks of frequencies in the three example cells after de-transformation, coloured in the same manner as panel (**a**). **e**, The scatter plot shows the relationship between the mean expression (x-axis) and variance (y-axis) for each gene after de-transformation. **f**, The scatter plot shows the distribution between the number of genes and the target sum (sum of all expression values) in the cell after de-transformation. **g**, The scatter plot shows the distribution between expression values and their ranks of frequencies in the three example cells after unscaling, displaying the ‘count-rank’ distribution. **h**, The dot plot shows the distribution between the mean expression (x-axis) with the variance (y-axis) for each gene in the count matrix. **i**, The scatter plot shows the distribution between the count values and the rounding errors after denormalisation. Each dot represents a count value in a cell and its rounding error.

### scDenorm recovers raw count matrices for large-scale database

To evaluate the performance of scDenorm in real scenarios, 40 processed datasets (**Supplementary Table 1**) from the UCSC Cell Browser^24^ were used as test data covering a good variety of species, tissues and sequencing techniques (**Fig. 3b**). Denormalisation performance was evaluated by two metrics: 1) rounding error, defined as the difference between a value in the denormalised matrix and its nearest integer (round value), and 2) recovery error, defined as the difference between a value in the normalised matrix and its corresponding value in the denormalised matrix after renormalisation (**Fig. 3a**, See **Methods**). 32 out of the 40 test sets were successfully denormalised (**Fig. 3c, Supplementary Table 1**), while the 8 unsuccessful cases were normalised as TPM (Transcripts per Million), log2(FRPM) or by scTransform^32^ method rather than the delta method (**Supplementary Table 1**). The mean versus variance distribution confirms a negative binomial distribution after denormalisation, indicating the successful denormalisation (**Supplementary Fig. 5**). The rounding errors, which are positively correlated to the expression value (**Supplementary Fig. 6**), are consistently below 0.005 (**Fig. 3e**). For recovery error, the absolute values are below 10^−6^ in 27 datasets normalised with natural logarithmic transformation (**Fig. 3d, Supplementary Table 1**), indicating a good accuracy of scDenorm (**Fig. 3f**). Further benchmarking of the denormalisation on 60 datasets (**Supplementary Table 2**) from the Brain Cell Atlas^30^ show similar results (**Supplementary Fig. 7**). In addition, scDenorm shows a linear computational time complexity and memory usage with increasing number of cells and genes, demonstrating a high computational efficiency and scalability (**Supplementary Fig. 4c,d**).

**Fig. 3.**
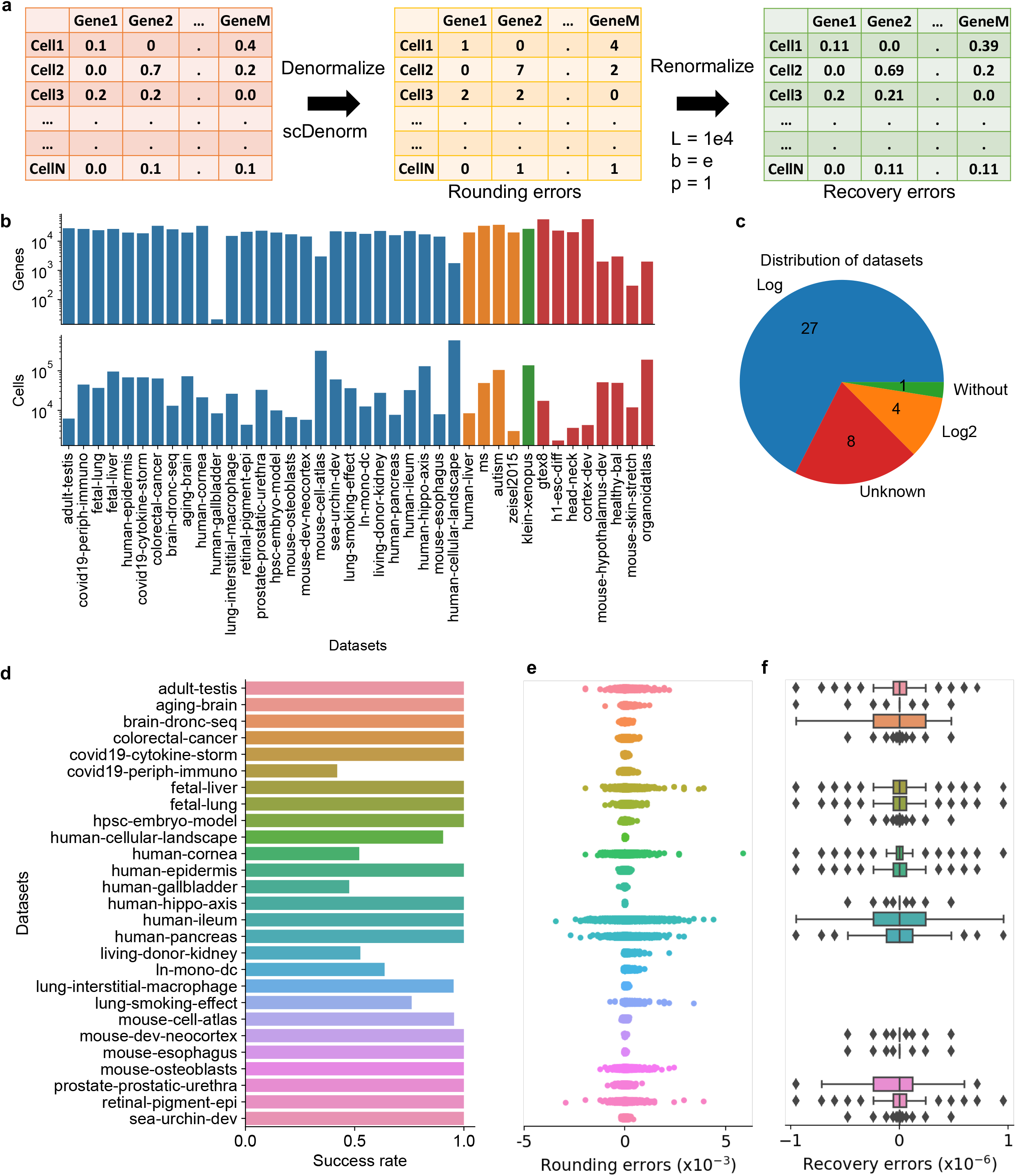
Performance of scDenorm on normalised scRNA-seq data from UCSC Cell Browser **a**, The diagram illustrates the workflow of scDenorm to evaluate denormalisation using the rounding and recovery error. The processed matrix (left) deposited in the UCSC database is first denormalised (middle) with scDenorm to calculate the rounding error (see **Methods**). Subsequently, the denormalised matrix (middle) is re-normalised (right) to measure the recovery error (see **Methods**). The values of target sum and pseudo count normalisation parameters are 1e4 and 1, respectively. **b**, Two barplots show the number of genes (top) and number of cells (bottom) for the collected UCSC datasets. The x-axis shows the datasets by name, while the y-axis shows the log-scaled number of genes (top) and the log-scaled number of cells (bottom). The colours represent the different parameters of the delta normalisation. Blue and orange are natural base (e) and base=2 respectively, green represents data without log-transformation. Red shows non-delta method normalisation cases, which could not be denormalised by scDenorm. **c**, The pie chart shows the distribution of the number of datasets classified by different base values, detected by scDenorm. The colours are the same as shown in panel (**b**). Unknown represents non-delta method normalisation cases. **d**, The bar plot shows the distribution of the success rate (see **Methods**) across the 27 datasets that were normalised with natural logarithmic transformation. **e**, The jitter plot shows the distribution of rounding errors observed in the denormalised datasets. The x-axis is the rounding error, while the y-axis shows the same datasets as panel (**d**). **f**, The box plot shows the distribution of recovery errors after re-normalization with the parameters of target sum, pseudo count and logarithm base as 1e4, 1 and natural base (e), respectively. The x-axis is the recovery error, while the y-axis shows the datasets in the same order as in panel (**d**).

### scDenorm accurately recovers raw counts in different scenarios

In realistic scenarios, denormalising the normalised matrix deposited in the database can be affected by several aspects, including: 1) the parameters used in delta normalisation method; 2) the digital precision kept in the deposited data; and 3) the genes filtered after data normalisation (**Fig. 4a**), (e.g., some lowly expressed genes could be removed). Using the 10x 3k PBMC single-cell dataset as a showcase, we benchmark these aspects.

**Fig. 4.**
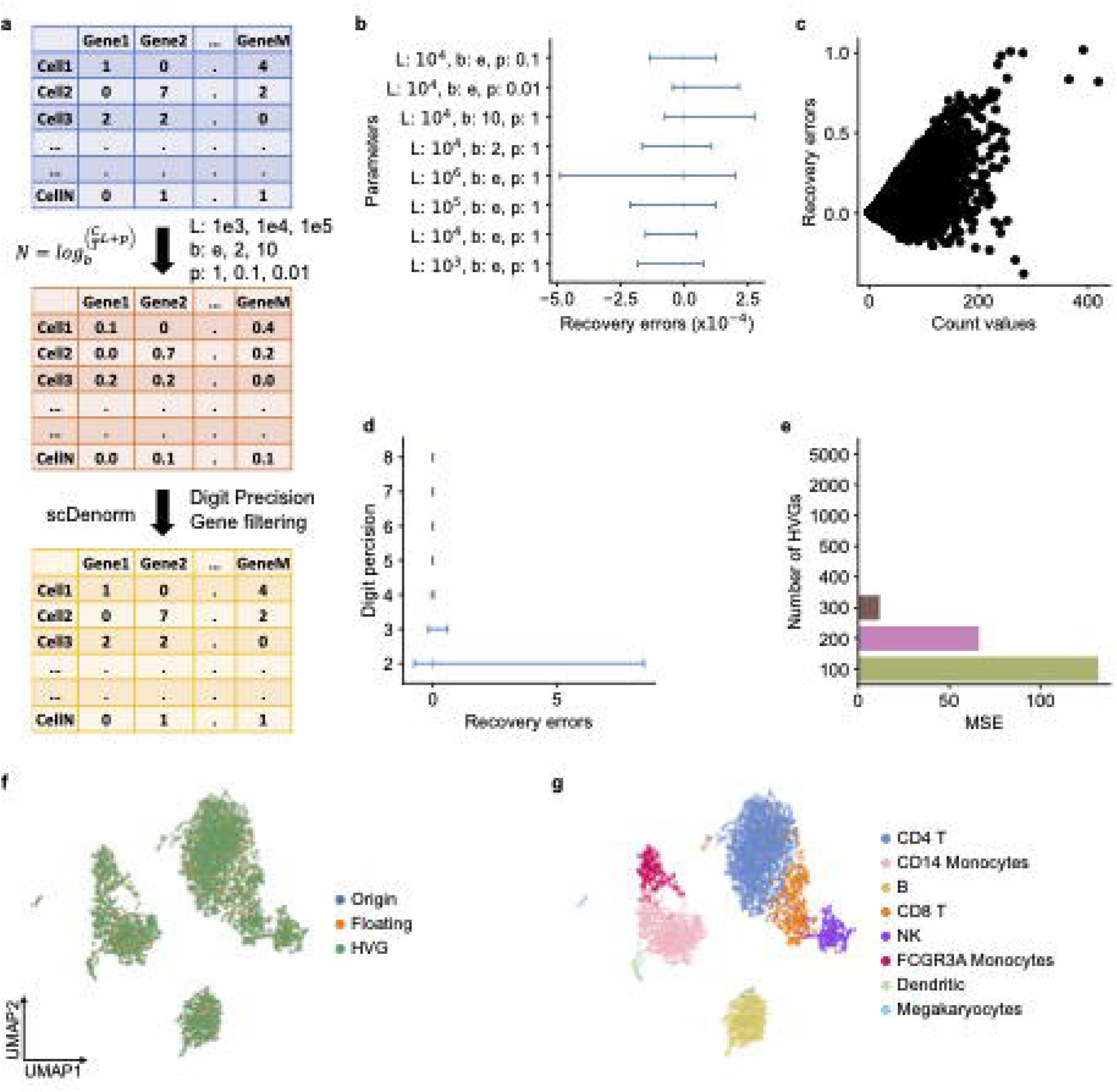
Benchmark of scDenorm on different normalisation parameters, digital precision and gene filtering **a**, The diagram shows the workflow of evaluating the recovery errors of denormalisation in three scenarios, different normalisation parameters, different digit precision, and gene filtering, on the PBMC 3k dataset. The raw count matrix (top) is first normalised with different parameter sets (middle), and then denormalised (bottom) with scDenorm giving different digit precisions and filtered genes. In delta method normalisation: C is the count value; T is the total count value of a cell; L is the target sum; p is the pseudo count; b is the base of logarithmic function; and N is the normalised gene expression values. **b**, The line plot shows the distribution of recovery errors from using different parameter sets in delta normalisation. **c**, The dot plot shows the distribution between raw count values and their recovery errors after the conversion of normalised data from float32 to float16. **d**, The line plot shows the distribution of recovery errors from normalised float32 data while preserving different levels of digit precision (2-8 digit precisions). **e**, The histogram shows the mean square error of regression loss of equation (4) (see **Methods**) from normalised data with different numbers of highly variable genes (from 100 to 5,000). **f**, The UMAP plot shows the distribution of data processed using different approaches, including original processed data (blue), denormalised data after converting to float16 (orange), and denormalised data after selecting 2,000 highly variable genes (green). **g**, The UMAP plot shows the cell type distribution of panel (**f**), colour-coded by cell types.

We examined the effect of normalisation parameters (target sum, log-transformation base, and pseudo count) by simulating the normalisation process with eight sets of hierarchical parameters. The dataset was normalised using these parameters, and denormalised by scDenorm. As shown in **Fig. 4b**, the errors between the denormalised value and its raw count in all denormalised matrices are consistently low, as < 5×10^−4^, indicating a minimal impact from normalisation parameters.

The digital precision of the normalised data, which can vary depending on the data processing tools and the saved file format, can also affect computational memory consumption. By default, the normalised data is saved as float32 (single-precision floating-point) format, with a precision of 6 to 9 decimal digits^33^. We simulated data with lower precision and denormalised it with scDenorm. The recovery error was less than 0.5 for count values less than 100, and less than one for count values greater than 100 (**Fig. 4c**). The errors are less than 0.01 when the digit precisions are more than 4 digits. The precision achieved with 3 to 4 decimal digits was consistent with the results of float16 conversion (**Fig. 4d**). Yet two decimal precision shows larger errors in highly expressing genes but keeps the cell identities (**Supplementary Fig. 8**).

In scRNA-seq data analysis, some genes expressed in few cells need to be removed or only selected genes may be kept in the normalised matrix. We simulated a gradient of the number of selected genes and tested the impact on denormalisation. No detectable error was found when more than 300 genes are kept in the normalised matrix, with the error increasing as the number of genes decreases (**Fig. 4e, Supplementary Fig. 9**). However, downstream data visualisation demonstrates that the denormalised matrices from float16 precision and a selection of 2000 highly variable genes successfully recovered the UMAP representation derived from raw counts (**Fig. 4f,g**), in spite of minor differences in the values.

### scDenorm facilitates downstream analysis

We further evaluated the impact of denormalisation on downstream analysis tasks, including data integration, cell type annotation, differential expression (DE) analysis, and gene ontology (GO) analysis. As data from different batches may go through different normalisation, three datasets were prepared to cover different batch types. The batch in the COVID-19 PBMCs^34^ dataset are samples from two patients, while in the human prefrontal cortex (PFC)^31,35^ dataset are samples from two different studies. And the batch in the human skin^36^ dataset are groups of samples from young and old donors.

The two patient samples in the COVID-19 PBMCs^34^ dataset are first normalised with different target sums (1000 and 10,000) before going through downstream analysis (**Supplementary Fig. 10**). Firstly, if without denormalisation, the UMAP visualisation after harmony integration shows cells of the same cell type exist in multiple clusters, e.g., plasmablast (**Fig. 5a**). Subsequently, CellTypist^37^, a well-established reference-based machine learning algorithm, was used to annotate the cell types. The first sample was used as a reference to annotate the cell types in the second sample, resulting in an accuracy of 66% (the consistency between the original cell type labels and those assigned by CellTypist). Notably, CD14+ monocytes were misclassified as plasmacytoid dendritic cells (pDCs) and hematopoietic stem and progenitor cells (HSPCs), while CD8+ T cells were misclassified as natural killer (NK) cells and CD4+ T cells (**Fig. 5b**). Fortunately, with the help of scDenorm denormalisation, the two patient samples can be integrated, with each cell type cluster forming distinct clusters (**Fig. 5c**). And the accuracy of cell type annotation using CellTypist increased to 92%, indicating effective correction of denormalisation. Furthermore, mis-annotated cell type labels may result in biases in differential gene expression analysis (**Fig. 5d, Supplementary Fig. 10d**) and gene ontology analysis (**Supplementary Fig. 10e**). For instance, 350 differential genes in HSPCs matched the ‘gold standard’ after scDenorm, compared to only 81 before scDenorm (**Fig. 5e**). Gene enrichment analysis of the differential genes in HSPCs indicated that the GO terms enriched after scDenorm closely aligned with those of the ‘gold standard’, whereas the enrichment before scDenorm showed minimal overlap (**Supplementary Fig. 10f**). The enriched GO terms are relevant functions associated with HSPC cells, such as hematopoietic stem cell proliferation and hematopoietic progenitor cell differentiation (**Fig. 5f**).

**Fig. 5.**
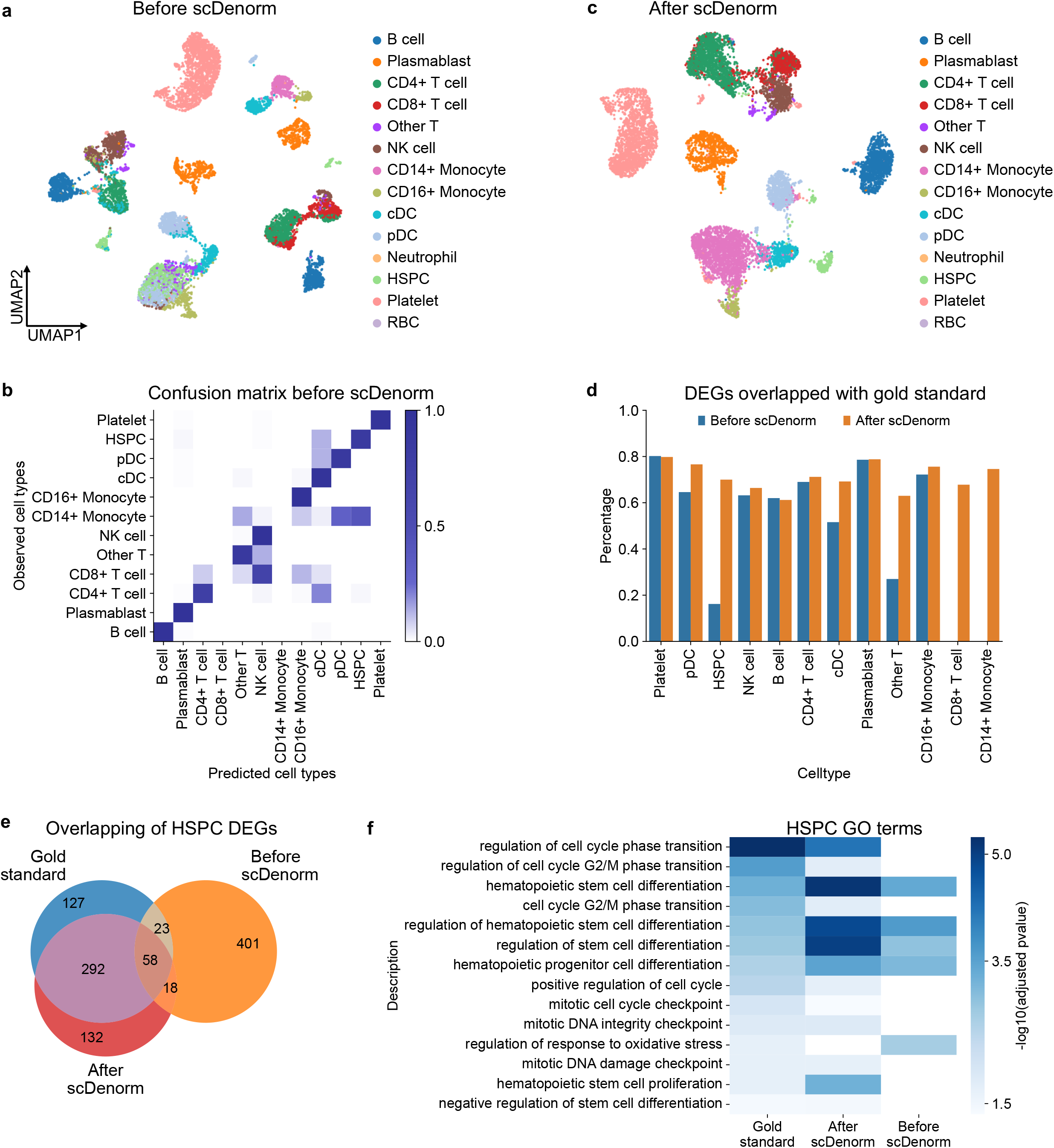
Different normalisations impact cell type annotation of COVID-19 PBMCs **a**, The UMAP plot shows the distribution of cells before denormalisation by scDenorm, coloured by CellTypist-predicted cell type annotation (see **Methods**). **b**, The UMAP plot shows the distribution of cells after Harmony data integration following by scDenorm. Cells are coloured by CellTypist-predicted cell type annotation. **c**. The heatmap shows the confusion matrix between the published cell type labels and the CellTypist-predicted cell types based on the Harmony-integrated latent space before scDenorm denormalisation. The x-axis represents predicted cell types, while the y-axis denotes original cell type annotation published in the study. **d**.The histogram shows the percentage of DEGs overlap between the ‘gold standard’ (DEGs derived according to the original published cell type labels) and the ones derived from re-analysis before (blue) and after (orange) scDenorm across cell types. The DEGs are calculated with a two-sided Wilcoxon test using the CellTypist-predicted cell types as clusters. **e**.The Venn diagram shows the overlap of the top 500 DEGs for HSPCs derived from the ‘gold standard’ (blue) from original study, as well as before (orange) and after (red) scDenorm. **f**.The heatmap shows the enriched Gene Ontology (GO) terms of HSPCs derived from DEGs of the ‘gold standard’ (blue) from original study, as well as before (orange) and after (red) scDenorm.

Similarly, the same analysis of data integration and cell type annotation was performed on two other datasets, the human prefrontal cortex dataset and the human skin dataset. Both study-wise batch (the former dataset) and condition-wise batch (the latter dataset) demonstrated that data processed by scDenorm yielded superior integration results (**Fig. 6, Supplementary Fig. 11**) and improved annotation results from CellTypist (**Supplementary Fig. 11g,h, Supplementary Fig. 12d,e**). Yet, the mislabelled cell types lead to biased differentially expressed genes (DEGs) (**Supplementary Fig. 11i**) and Gene Ontology (GO) terms (**Supplementary Fig. 11j**).

**Fig. 6.**
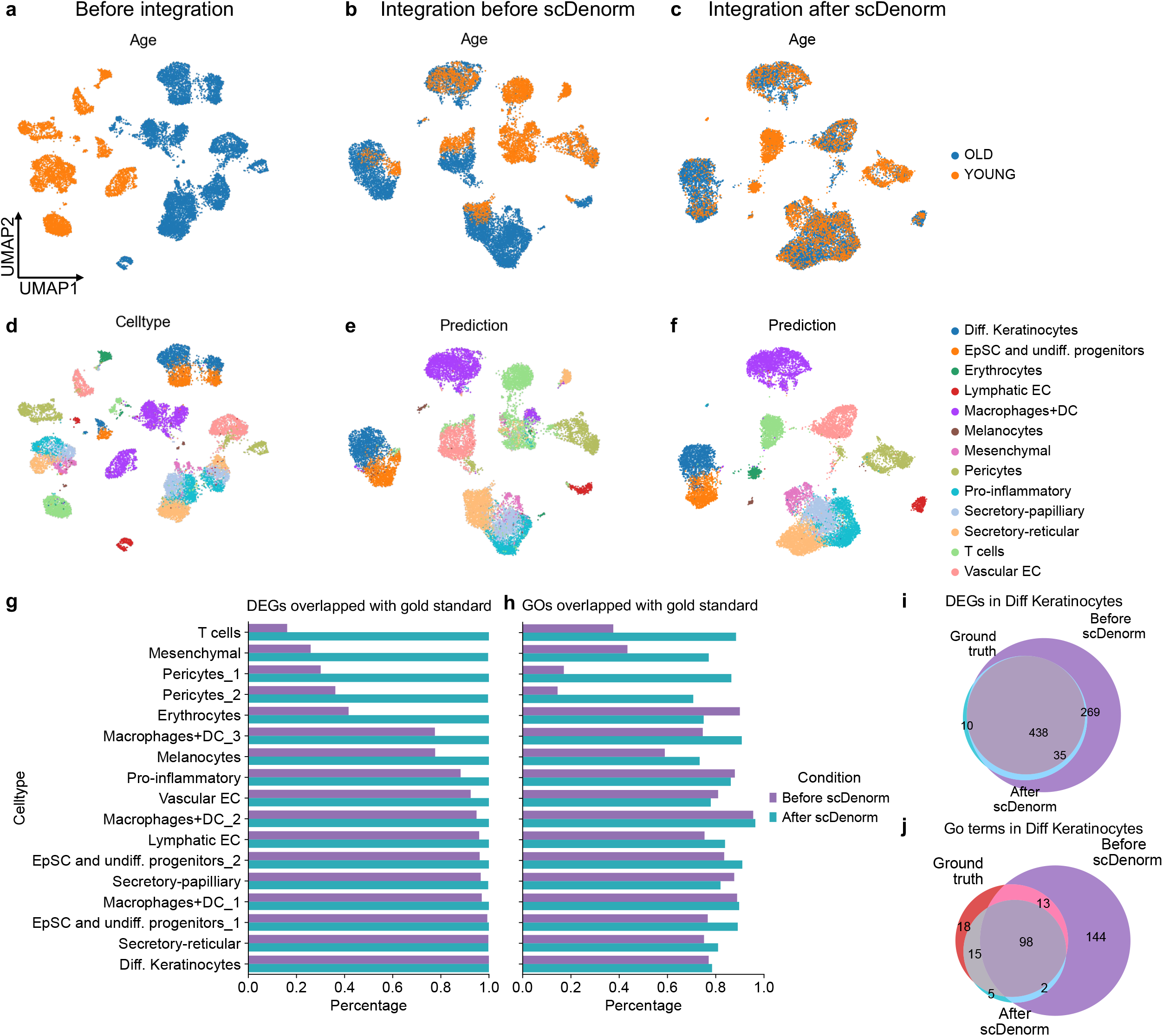
Different normalisations impact differential expression analysis **a**, The UMAP plot shows the cell distribution of the two age groups of the human skin dataset (Solé-Boldo et al.) before data integration without the scDenorm denormalisation, coloured by age groups. **b**, The UMAP plot shows the Harmony-integrated result without the scDenorm denormalisation, coloured by age groups. **c**, The UMAP plot shows the Harmony-integrated result after running scDenorm, coloured by age groups. **d**, The UMAP plots show the same distribution as panels (**a**), coloured by original cell type labels from Solé-Boldo et al. **e-f**, The UMAP plots show the same distribution as panels (**b**-**c**), but coloured by CellTypist-predicted cell type labels before and after denormalisation by scDenorm. **g-h**, The histograms show the percentages of DEGs (**g**) and GO terms (**h**) overlap between the ‘gold standard’ (DEGs extracted from the data in the original study, GO terms derived from these DEGs) and data before (purple) and after (blue) scDenorm across cell types, using the ‘gold standard’ cell type labels reported in the original publication. The DEGs are calculated with a two-sided Wilcoxon test based on the original cell type from the human skin dataset (Solé-Boldo et al.), while the GO analysis shows Benjamini-Hochberg-adjusted P value <0.05. **i-j**, The Venn diagrams show the overlaps of the DEGs (**i**) (Benjamini-Hochberg-adjusted P value <0.05 and logFC >0.25) and GO terms (**j**) for differentiated keratinocytes derived from data of the ‘gold standard’ (red), as well as before (purle) and after (blue) scDenorm denormalisation.

However, the correct cell type annotation can also be achieved according to marker genes. We evaluate the impact of normalisation parameters on downstream DE and GO analysis. Taking the human skin data^36^ as an example, differential gene expression analysis was performed before and after scDenorm using the cell type labels derived from the original publication, while the published differential expression genes (see **Methods**) and their resulted GO results were taken as ‘gold standard’. The DE and GO results after scDenorm show higher consistency with the ‘gold standard’ than the results before scDenorm (**Fig. 6g,h**). In addition, the DEGs identified before scDenorm include more false positive genes (**Fig. 6i**), resulting in the enrichment of unrelated GO terms (**Fig. 6j**), such as the nuclear transport function for keratinocyte cells (**Supplementary Fig. 12f**).

## Discussion

In our survey of 133 well-established single-cell studies, delta method normalisation takes up >83% (110) of the datasets (**Supplementary Table 3**), since it is implemented in widely-used SCANPY and Seurat analysis workflows. We demonstrate the capability of scDenorm on an example dataset and large-scale test sets from UCSC database^24^ and the Brain Cell Atlas^30^. Different parameter sets in the delta method normalisation as well as the digital precision kept in the normalised data have minimal effect on denormalisation. Moreover, the number of genes kept after normalisation does not significantly affect denormalisation, unless the number of genes used is too small (less than 300) (**Fig. 4e**). In the 40 datasets from UCSC database (**Supplementary Table 1**) and 60 datasets from Brain Cell Atlas (**Supplementary Table 2**), scDenorm successfully restored count values in the majority (88%) of the cases, with minimal rounding errors and recovery errors. Therefore, scDenorm may robustly recover matrices for the majority (estimated to be 80-90%) of the datasets, which are delta method normalised, while maintaining efficient computational speed (**Supplementary Fig. 4c**).

The limitations of scDenorm relies on specific prerequisites of the delta method normalisation. Datasets normalised using alternative methods may not be compatible with scDenorm. For example, GLM residual methods (such as SCTransfrom^32^) and latent expression (such as Sanity^38^ and Dino^39^) cannot be denormalised by scDenorm. Fortunately, other denormalisation methods besides the delta method only constitute 10-20% of the datasets. And the raw counts of these datasets can be obtained from reads mapping. Additionally, the performance of scDenorm may be influenced by the choice of normalisation parameters and the quality of the input data. Cells whose gene expression distribution deviates from the assumptions of negative binomial distribution may lead to the failure of the denormalisation process.

Several case studies show that different normalisations can result in unnecessary deviations in downstream analysis, including data integration, cell type annotation, differential gene expression, gene ontology and pathway analysis. In particular, biased differential expression or gene ontology results can be generated due to different normalisation parameters when the cell type annotation is correct. Therefore, denormalising the expression matrix to raw counts can be a good choice to mitigate biases in downstream analysis. It could be a key question for large-scale data integration, where study-wise batch effects need to be minimised while biology should be kept. Batch correction as well as data integration methods have already been extensively discussed and benchmarked^8^. Here we highlight the consistency in data processing, which is nontrivial when data from tens or hundreds of studies need to be combined together. Consistent single-cell data analysis workflows which can both keep the raw conclusions from the publications and get integrated with data from other studies would greatly help. Therefore, the availability and reproducibility of the raw published analysis code would be important.

## Conclusions

Here, we demonstrate that inconsistent data normalisation can generate unexpected bias in data integration, potentially obstructing atlas-level single-cell data integration. Fortunately, denormalising processed data back to raw counts could standardise analysis, thereby facilitating the creation of comprehensive cell atlases. We present scDenorm, a tool designed to denormalise data from the delta method normalisation, which are widely used by 80% of the 40 datasets in the UCSC database and 93% of the 60 datasets in Brain Cell Atlas. It employs both equation solving and regression methods to determine the parameters in the delta method. Benchmarks on 32 UCSC cell browser datasets and 56 Brain Cell Atlas datasets demonstrate the efficacy of scDenorm for delta method normalisation data, with further applications on COVID-19 PBMCs, prefrontal cortex, and human skin datasets revealing its ability to mitigate biases in downstream analysis. scDenorm can be a useful tool in atlas level single-cell data processing and integration, such as the Human Cell Atlas^40^, The Human Developmental Cell Atlas^41^, the Brain Cell Atlas^30^, and HuBMAP^42^.

## Methods

### Assumption and algorithm design

In scRNA-seq, the data is in the form of a count matrix, where most entries are zeros due to the sparsity of gene expression. Our assumption is that the scRNA-seq data follows negative binomial distribution, which is theoretically and empirically well-supported for the unique molecular identifier data^17^. This means that probabilistically speaking, in the count matrix zero is the most frequently observed count, followed by one and two and so on. The sequential pattern of these values has a probabilistic one-to-one correspondence with the rank of their frequency by descending order (**Fig. 1f**). The smaller the values, the higher the probability of the correspondence (**Fig. 1g**). For example, without considering 0, the probability that the values 1 and 2 are equal to the rank of their frequency is almost 100%. Based on this assumption, we design an algorithm to normalise scRNA-seq data that has been normalised by the most commonly used delta methods, which scale the raw counts by the total number of counts (library size) and target sum (the summed value of the cell after scaling), and then log-transformed after adding a pseudo count (**Supplementary Fig. 3a**). Specifically, we consider a scaled expression matrix from a count matrix which has been transformed to adjust for differences in the scale of the features (e.g., genes) in the data. In scRNA-seq data, a scaled expression matrix typically refers to a count matrix that has been normalised and transformed to have a similar distribution of gene expression values across cells. For example, scRNA-seq data can be normalised to account for differences in sequencing depth and other technical factors that can affect the distribution of counts across cells and genes, such as total count normalisation. And it can be transformed to adjust for the distribution of gene expression values across cells such as log-transformation, and variance-stabilising transformation. The normalised gene expression matrix is derived from the count matrix to adjust for differences in gene expression across cells, which usually involves scaling and transformation techniques such as total count scaling and log-transformation. This normalisation process does not change the one-to-one correspondence between the gene expression value and its rank of the value’s frequency in a cell.

Using the probabilistic one-to-one correspondence property, we can extract a cell vector from a normalised expression matrix and sort the values based on their frequency in the vector. This allows us to establish that the most frequently occurring non-zero value corresponds to one and the second most frequent represents two and so forth, which means the rank number and the count number are theoretically the same and this is normally true for the top ranks. By following this procedure, we were able to obtain the rank and normalised value pairs (C, N) (where C is the rank and N is the normalised count) for the equation 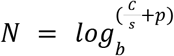, (s is the scaling factor, b is the base of log-transformation, and p is the pseudo count). First, we try reversing log transformation of natural base(e), base 2 and base 10, and solve the equation on the pairs of values (1, N_1_) and (2, N_2_), N_1_ and N_2_ are the values of the two most frequent numbers. Normally, we think the pseudo count C is given as 1. Otherwise, we need to check whether the variance of the solved C from different cells is sufficiently small, since each vector from the gene expression matrix has been augmented with the same pseudo count. If the unscaling process is unsuccessful for all of the above cases, we conclude that the matrix has not been pre-processed according to the workflow. The following shows the complete workflow of scDenorm algorithm.

The denormalisation algorithm can be divided into two steps, de-transformation and unscaling. In de-transformation, there are two sequential steps. Firstly, (a) we search for empirical values for the log-transformation bases and the pseudo count. It searches for empirical bases such as 2, e (natural base), and 10, as well as common pseudo counts like 0, 0.01, 0.1, and 1. If the pseudo count is 0, it indicates that the normalisation process has not added the pseudo count. A fraction of cells is used to evaluate if any of these bases or pseudo counts meet the criteria in step (c). if passing the criteria, skip to step 2. Otherwise, it goes to step (b) to determine the parameters. Step (b) uses equation solving method to determine the parameters: This method uses the two values (*N*_1_,*N*_2_) occuring most frequently in a cell to construct the following equation. For each cell i:

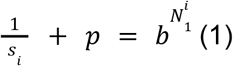

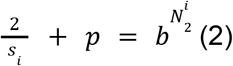

*s*_*i*_ is the scaling factor for the cell i. The p and b are pseudo count and base. From equation (1) and (2), we can get equation (3).

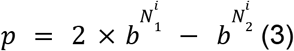

Randomly select a group of cells (e.g., n=100) to generate a corresponding set of data points (*N*_1_,*N*_2_), and solve p and b by equation (4) with optimization methods.

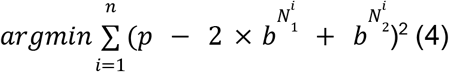

The L-BFGS-B method from the sklearn^43^ package is used to find the best base (b) and pseudo count (p). This method is based on the limited-memory Broyden-Fletcher-Goldfarb-Shanno (BFGS) algorithm, which is capable of large-scale optimization. L-BFGS-B allows for box constraints, ensuring that the parameters stay within specified bounds during optimization.

After the de-transformation, the sum of each cell should be the same or very similar. Step (c). checks if the sum of each cell is the same. For example, let X as the vector of the sums, x is a number in it. And if abs(x-mean(X)) is always smaller than the small number (*e*.*g*., mean(X)=10000, x=9999.7, small number is 0.5), then the de-transformation is successful. However, this is an ideal situation. Often, we encounter that after normalisation, the data filters out some genes for the quality control in downstream analysis. In addition, some normalisation methods do not scale the total expression values to the same for all cells. To address these complex cases, we also added the following criteria. If it is the automatic detection method, we only need to make sure that the mean square error (MSE) is small enough, such as 10^−5^. In general, we just need to unscale a cell to see if it is successful.

In unscaling, we have two approaches implemented in the same function, while a parameter can be used to select the option. The first approach (a) is based on regression to determine the scaling factors for all cells. The scaling factor is derived from fitting a regression model to the relationship between the de-transformed values and their ranks, providing an estimate of the scaling factor for each cell. For each cell:

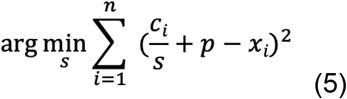

*c*_*i*_ is the rank, and *x*_*i*_ is the de-transformed value.

To ensure a more accurate one-to-one correspondence, only the first 5 pairs of values (*c*_*i*_, *x*_*i*_) are used. We can get the scaling factor s by optimising the equation (5) using the same L-BFGS-B method as in solving equation (4).

The second approach (b) is solving equations between the top two most frequent values: This method only used the first 2 pairs of values (*c*_*i*_, *x*_*i*_). We can get a closed form of solution by solving the following equation. For each cell:

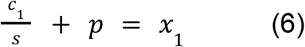

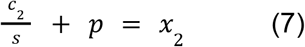

From equation (6) and (7), we can get equation (8).

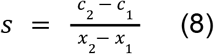

To evaluate the success of the denormalisation process, we measure the error between the denormalised value and its rounded value. Ideally, after denormalisation, the values should closely approximate integers. We can calculate abs(round(x)-x) and check if it is smaller than a certain number. Of note, in some cases, the same top value (e.g., 1) can be normalised into more than 1 different values due to some improper data processing, and the ranks of these numbers are thus lower than expected. These numbers with tiny differences are merged as one value by decreasing their digital precision.

scDenorm is publicly available as an open-source Python package and provides a user-friendly python function interface, which can be combined in the use of SCANPY analysis. It can be used both at the command line and interactively in Jupyter notebook. Description of the function details are provided in **Supplementary Materials**. Considering that different samples in a dataset may be normalised with different parameter sets, scDenorm also implements a per sample denormalisation function overloading the original ‘scdenorm’ function with a ‘by=sample’ parameter as input.

### Integration of scRNA-seq data from different normalisation parameters

We downloaded a PBMCs scRNA-seq data from the 10x genomics datasets (http://cf.10xgenomics.com/samples/cell-exp/1.1.0/pbmc3k/pbmc3k_filtered_gene_bc_matrices.tar.gz), and preprocessed and annotated the data according to this tutorial (https://scanpy-tutorials.readthedocs.io/en/latest/pbmc3k.html). Then, we used different parameter combinations (including 1e3, 1e4, 1e5, 1e6 as target sums, 2, e, 10 as base, and 1,0.1, 0.001 as pseudo counts) to normalise the data separately and merge all the data together. Principal Component Analysis (PCA) of 50 components was derived from the expression matrix. Three single-cell data integration tools (Harmony, BBKNN, scanorama) were tested to integrate the combined data with the normalisation parameters as the batch key. For data visualisation, Uniform Manifold Approximation and Projection (UMAP)^44^ is calculated in the integrated latent space or the PCA space.

### Consistency of count-rank relationship across sequencing platforms

The consistency of count-rank relationship refers to the percentage of the correct one-to-one correspondence between the gene expression value and its rank of the value’s frequency in a cell. For example, given 100 cells, we first calculate the frequency of the raw count values (the raw count value called as count) in each cell, and order the frequencies from highest to lowest. The order is called as rank, which ranges from 1,2, …, n. If count is the same as rank, we consider this to be a correct one-to-one correspondence. Finally, for the count from 1 to 10, we respectively calculate what percentage of cells have the correct one-to-one correspondence as the consistency of the count-rank relationship. To compare different sequencing platforms, we calculated the consistency of the count-rank relationship in 105 datasets obtained from the Brain Cell Atlas (http://braincellatlas.org). Among these datasets, 81 are from Chromium, 15 from Drop-seq, and 9 from Smart-seq2.

### Evaluation metrics

When benchmarking denormalisation for scRNA-seq data, two measures can be used: rounding error and recovery error. Rounding error measures the discrepancy between the denormalised values and their rounded counterparts. After denormalisation, the expected outcome is that the denormalised values approximate integers. Rounding error quantifies the extent to which the denormalised values deviate from integers. To calculate rounding error, the difference between each denormalised value and its rounded value is computed, see equation (9) below. Recovery error evaluates the difference before denormalisation and after re-normalising the denormalised values (values after scDenorm, **Fig. 3a**). To calculate recovery error, the difference between each normalised value and its re-normalised value is computed, see equation (10) below.

Specifically, we assume x is the normalised value (a single value for one gene in one cell), y is the denormalised value after scDenorm, and z is the re-normalised value from denormalised value (y). The rounding error is calculated as the difference between the denormalised value (y) and its rounded value, equation (9):

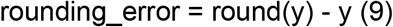

The recovery error is calculated as the difference between the normalised value (x) and the re-normalised value (z), equation (10):

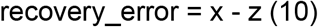

In certain cases, not all cells can be successfully denormalised due to poor sequencing quality, or a low number of genes. To evaluate denormalisation in such situations, we define success rate as the percentage of successfully denormalised cells, equation (11).

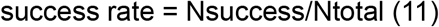

Nsuccess is the number of successfully denormalised cells, while Ntotal is the total number of cells.

### Benchmark scDenorm based on digital precision and gene filtering

To assess the impact of different digital precision of normalised data on the denormalisation process, we performed the following steps on the PBMC data: First, we applied total-count normalisation (the normalize_total function in SCANPY^19^) to the data matrix with target sum as 10,000, and log-transform (natural base, e) the data with 1 as pseudo count. Next, we used the round function to retain the data at different levels of precision, ranging from 2 to 8. Float16 corresponds to 3 to 4 decimal places of precision, while float32 corresponds to 6 to 9 decimal places of precision. Finally, we denormalise the data separately for each precision level and compare the results with rounding errors to evaluate their effects.

To test our algorithm for gene filtering on extreme cases, we selected a series of highly variable genes, ranging from 100, 200, 300, 400, 500, 1000, 2000 to 5000. Specifically, First, we normalise the data by sc.pp.normalise_total with target_sum as 10,000 and logarithmic the data with sc.pp.log1p. The high-variable genes are then selected using sc.pp.highly_variable_genes with layer as ‘count’ and flavour as ‘seruat_v3’. Finally, we use scDenorm to denormalise the data and calculate the recovery errors.

### Benchmark on large-scale datasets

To perform the usage of our algorithm on atlas data, we downloaded 40 datasets from the UCSC Cell Browser and 60 datasets from the Brain Cell Atlas, ensuring that they encompass a diverse range of species, sequencing platforms, and normalisation methods. First, we used scDenorm to denormalise each dataset. If successful, we calculate the rounding errors for the dataset, which quantifies the difference between the denormalised values before and after rounding. Additionally, if all total values (the sum of all the denormalised values in each cell) of the dataset are close to a fixed value (target sum, e.g., 10,000) after de-transformation, the recovery errors are calculated.

### Dataset processing for data integration and downstream analysis

The COVID-19 PBMC dataset from Arunachalam et al.^34^ (**Fig. 5**) was downloaded from GEO^45^ under the accession code GSE155673. Two samples Arunachalam_cov11 (S1) and Arunachalam_cov11 (S2) were processed with different delta normalisation parameters: S1 was normalised by target sum 1e3, while S2 was normalised by target sum 1e4. And both were log-transformed. For data visualisation, we performed Harmony^13^ data integration of these two samples after PCA of 50 components. For cell type annotation, CellTypist^37^ was used. S2 was used as the reference for annotating S1.

The human prefrontal cortex data includes datasets from two studies, Ma et al.^35^ (170,000 cells) and Velmeshev et al.^31^ (100, 000 cells) (**Supplementary Fig. 11**). The Velmeshev’s dataset was normalised to target sum 1e3 and logarithmic transformation, while the Ma’s dataset was not normalised. Harmony was used for data integration. For cell type annotation, the Ma’s dataset was used as the reference for annotating Velmeshev’s dataset.

The human skin dataset from Solé-Boldo et al., was downloaded from GEO under the accession code GSE130973 (**Fig. 6**), including two young (25 and 27 years old) and three old (53, 69 and 70 years old) donors. The young samples were normalised to target sum 1e3, while the old samples were normalised to target sum 1e4. And both samples were logarithmic transformed after normalisation. For cell type annotation, the old samples were used as the reference for annotating the young sample.

For data processing after denormalisation with scDenorm, we follow a standard workflow of data normalisation and dimension reduction. Specifically, the expression matrix is normalised to target sum of 10,000 and log-transformed. And the default dimension reduction process in the SCANPY workflow was used, including PCA, Harmony integration and UMAP visualisation. CellTypist was used to predict the cell types as described above.

Downstream analysis after cell type annotation includes differential expression (DE) analysis and Gene Ontology (GO) pathway analysis. Differential gene expression analysis was conducted for each cell type (one against the rest) using the Wilcoxon test implemented in SCANPY^19^. As part of the dataset is used as the reference dataset, the differentially expressed genes (DEGs) derived from the reference dataset is used as ‘gold standard’ in our evaluation. The same approach was used for the calculation of the DEGs before and after scDenorm. The top differentially expressed genes were compared across different thresholds (top 50, 100, 200, 500, and 1000). As for GO pathway analysis, the enrichGO program was used on the top 500 differential expression genes.

DE and GO analysis with the correct labels from Solé-Boldo et al. was conducted for each cell type (one against the rest) using the Wilcoxon test implemented in Seurat (V3.1.1), same as the version described in Solé-Boldo et al. These analyses were conducted before scDenorm and after scDenorm. The differentially expressed genes were obtained from the study’s supplementary materials as ‘gold standard’.

## Supporting information

Rebuttal

## Availability of Data and Materials

The 10x 3k PBMC data was downloaded from the 10x genomics website https://support.10xgenomics.com/single-cell-gene-expression/datasets/1.1.0/pbmc3k. The dataset with both the normalised expression and the raw count matrix was downloaded from https://autism.cells.ucsc.edu. 40 processed datasets (**Supplementary Table 1**) were downloaded from https://cells.ucsc.edu. 60 processed datasets (**Supplementary Table 2**) were downloaded from https://www.braincellatlas.org. Datasets used in the manuscript have been deposited at zenodo: https://doi.org/10.5281/zenodo.11240443. Codes for reproducing this work are available via GitHub at: https://github.com/rnacentre/scDenorm-reproducibility.

The source codes and documentation of scDenorm are available at https://github.com/rnacentre/scDenorm. It can also be installed via pip or anaconda, at: https://pypi.org/project/scDenorm/ and https://anaconda.org/changebio/scdenorm, respectively. This software is licensed under the Apache License 2.0, an open source licence compliant with Open Source Initiative.

## Declarations

### Ethics approval and consent to participate

Not applicable.

### Consent for publication

Not applicable.

### Competing interests

The author declares no competing interests.

### Author’s contributions

Z.M. and A.B. designed and conceived the study. Y.H., Z.M. and L.Y. implemented the scDenorm algorithm, Y.H. conceived and performed most of the bioinformatics analyses. Y.A. performed part of the analysis. H.Z., Y.Z., S.L., X.Y., and M.S. provided some datasets. Y.H., Z.M., A.V.P., Y.A., I.P. wrote the manuscript. Z.M., M.S., X.Y. supervise the study.

## Acknowledgements

The authors thank Ziliang Huang for help with the datasets.

## Funding

This work was supported by the National Key R&D Programs of China (2023YFF1204700, 2024YFF1206600), the Natural Science Foundation of China (32270707), the Major Project of Guangzhou National Laboratory (grant nos GZNL2024A01002 and GZNL2023A01006), the R&D Programs of Guangzhou National Laboratory (grant nos HWYQ23-003 and YW-YFYJ0102), and Postdoctoral Research Project Funding of Guangzhou, BSHF23-049.

## Notes

### Competing Interest Statement

The authors have declared no competing interest.

