## Supplementary information for "scDenorm: a denormalisation tool for integrating single-cell transcriptomics data"

[1] GMU-GIBH Joint School of Life Sciences, Guangzhou National Laboratory, Guangzhou Medical University, Guangzhou, China
[2] European Molecular Biology Laboratory, European Bioinformatics Institute, EMBL-EBI, Wellcome Genome Campus, Cambridge CB10 1SD, UK
[3] Department of Obstetrics and Gynaecology, Harbin Red Cross Central Hospital, Harbin 150001, China.
[4] Department of Surgery Oncology, Harbin Medical University Cancer Hospital, Harbin 150001, China
[5] State Key Laboratory of Ophthalmology, Zhongshan Ophthalmic Center, Sun Yat-sen University, Guangdong Provincial Key Laboratory of Ophthalmology and Visual Science, Guangzhou, China
[6] Guangdong Province Key Laboratory of Brain Function and Disease, Guangzhou, China
[7] School of Life Science and Technology, Harbin Institute of Technology, Harbin 150001, China
[8] Translational Research Institute of Brain and Brain-Like Intelligence and Department of Anesthesiology, Shanghai Fourth People's Hospital Affiliated to Tongji University School of Medicine, Shanghai, China
* to whom correspondence should be addressed

#### Supplementary Figures

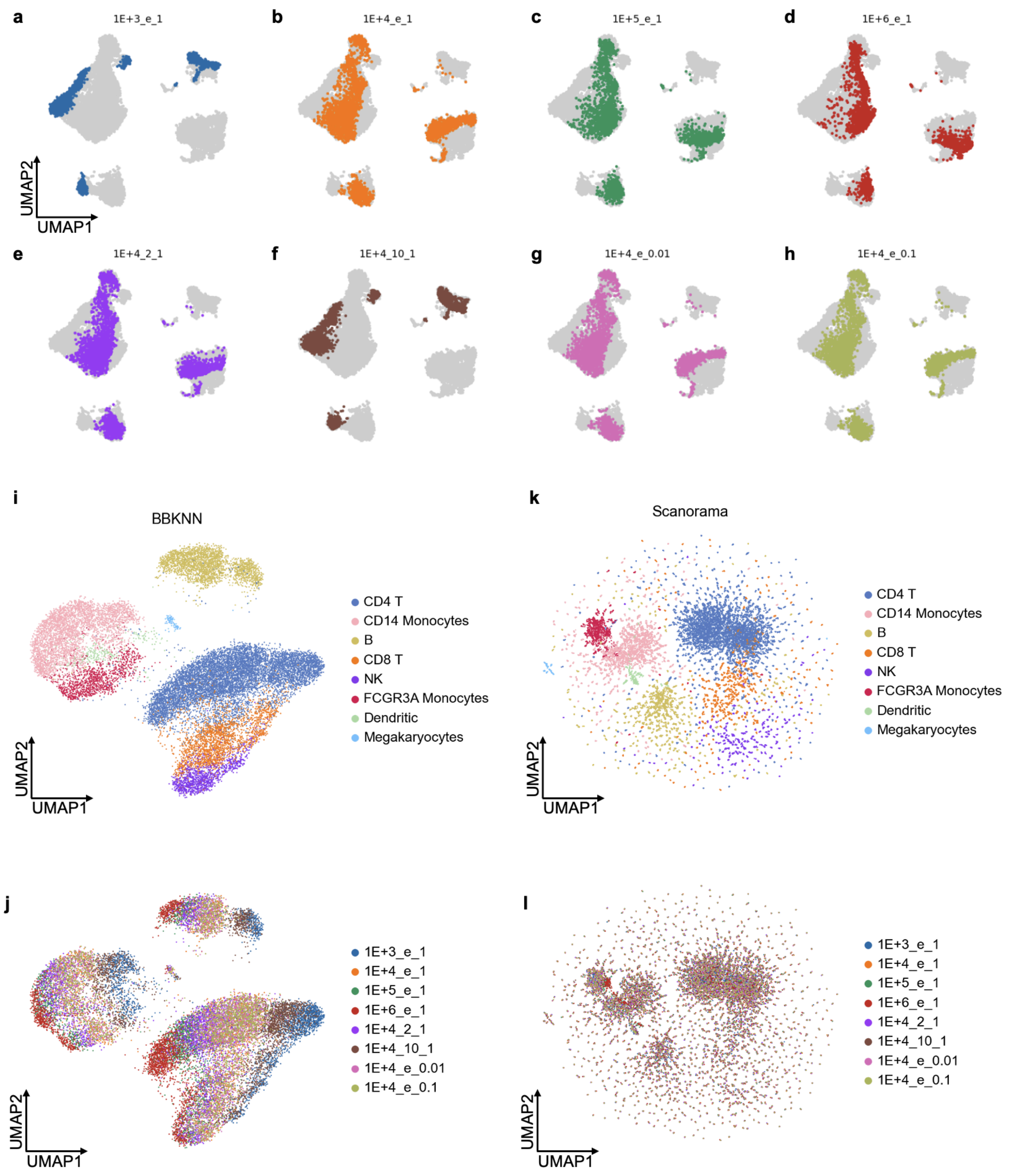


Supplementary Fig. 1 Inconsistent data normalisation generates bias in data integration

**a-h**, Each plot shows the UMAP based on PCA space from data obtained with different normalization methods, including different combinations of library size (L), base (b) and pseudo count (p).

**i-j**, UMAP plots the BBKNN-integrated result on the data obtained from different normalization methods after BBKNN integration, coloured by different cell types (**i**) and different normalization methods (**j**).

**k-l**, UMAP plots show the scanorama-integrated result on the data obtained from different normalization methods after Scanorama integration, coloured by different cell types (**k**) and different normalization methods (**l**).

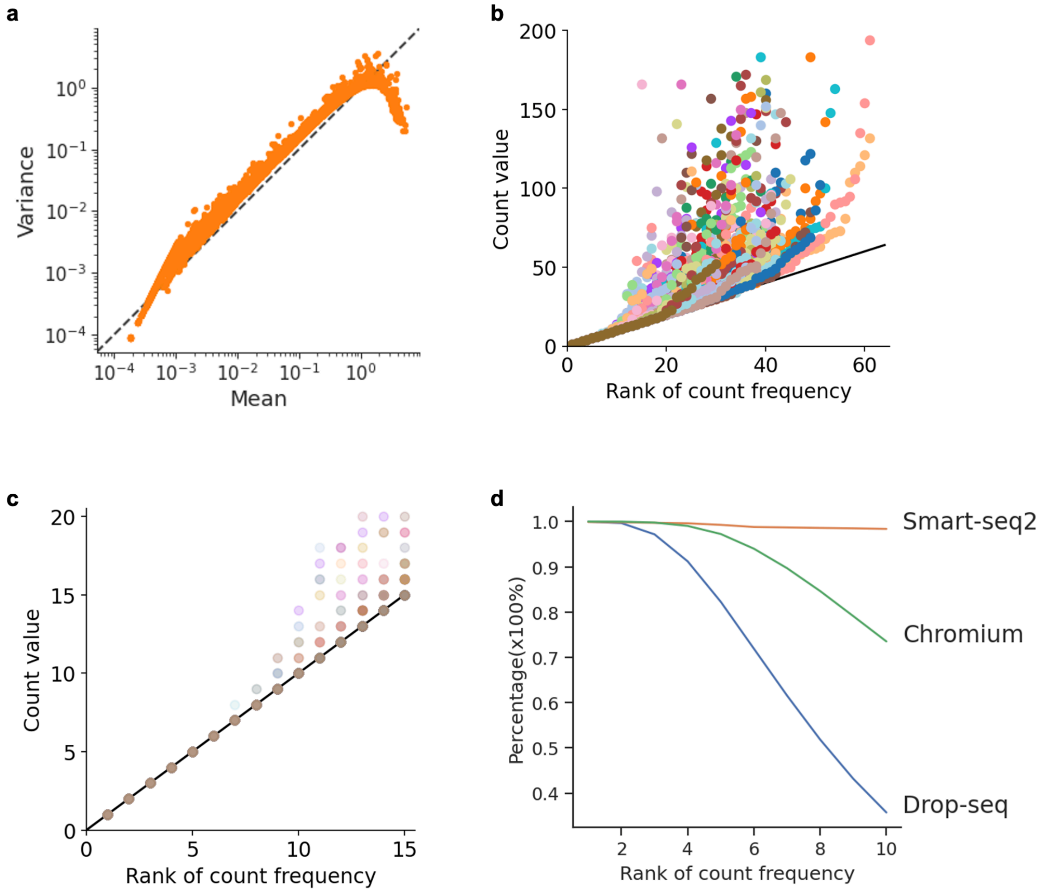


### Supplementary Fig. 2 Features of droplet-based single-cell data

**a**, The relationship between the log-transformed mean expression (x-axis) with the log-transformed variance (y-axis) for each gene in the count matrix from the study (Satija et al., 2015).

**b**, The relationship between the rank of the count frequency (x-axis) with the count value (y-axis) in cells.

**c**, the zoom-in view of panel (b).

**d**, The change of the percentage of cells with the value of count equal to its rank in three scRNA-seq technologies (Chromium, Smart-seq2, and Drop-seq).

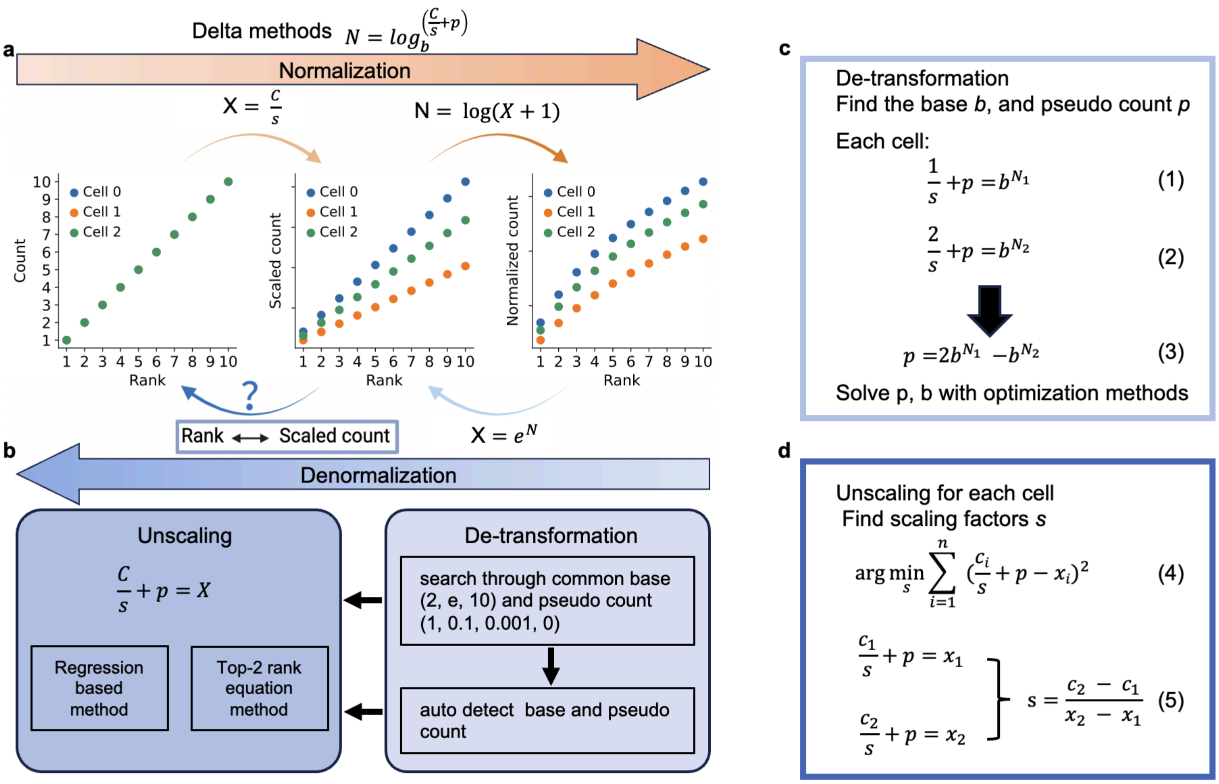


#### Supplementary Fig. 3 Schematic representation of scDenorm algorithm

**a,** Standard pre-processing: routinely droplet-based single-cell data are first normalised by a size factor (s, usually total count with a library size factor) with and pseudo count (p) secondly transformed by a logarithmic function with a base (b) for downstream analysis.
**b,** Denormalisation: scDenorm performs the reverse of the task above. First it reverses the logarithmic transformation by searching for common bases (e.g., 2, e, 10) and common pseudo counts (e.g., 0.01, 0.1, 1), or auto detect base and pseudo count. Then, it reverses the scaling by determining the scaling factor for each cell.
**c**, The formulas and derivations of automatic detection method for base and pseudo count during the de-transformation process. b is the base, p is the pseudo count, s is the scaling factor, and $N_1$ and $N_2$ are the most and second frequency values for each cell.
**d**, The formulas of solving the scaling factor for each cell during the unscaling process. Equation (4) is the formula of the regression based method, and equation (5) is the formula of the top-2

rank equation method. $c_i$ are the ranks of the frequency of normalised values ( $c_1$ is 1, and $c_2$ is 2, etc). $x_i$ are the values sorted by frequency in a decreasing order after de-transformation.

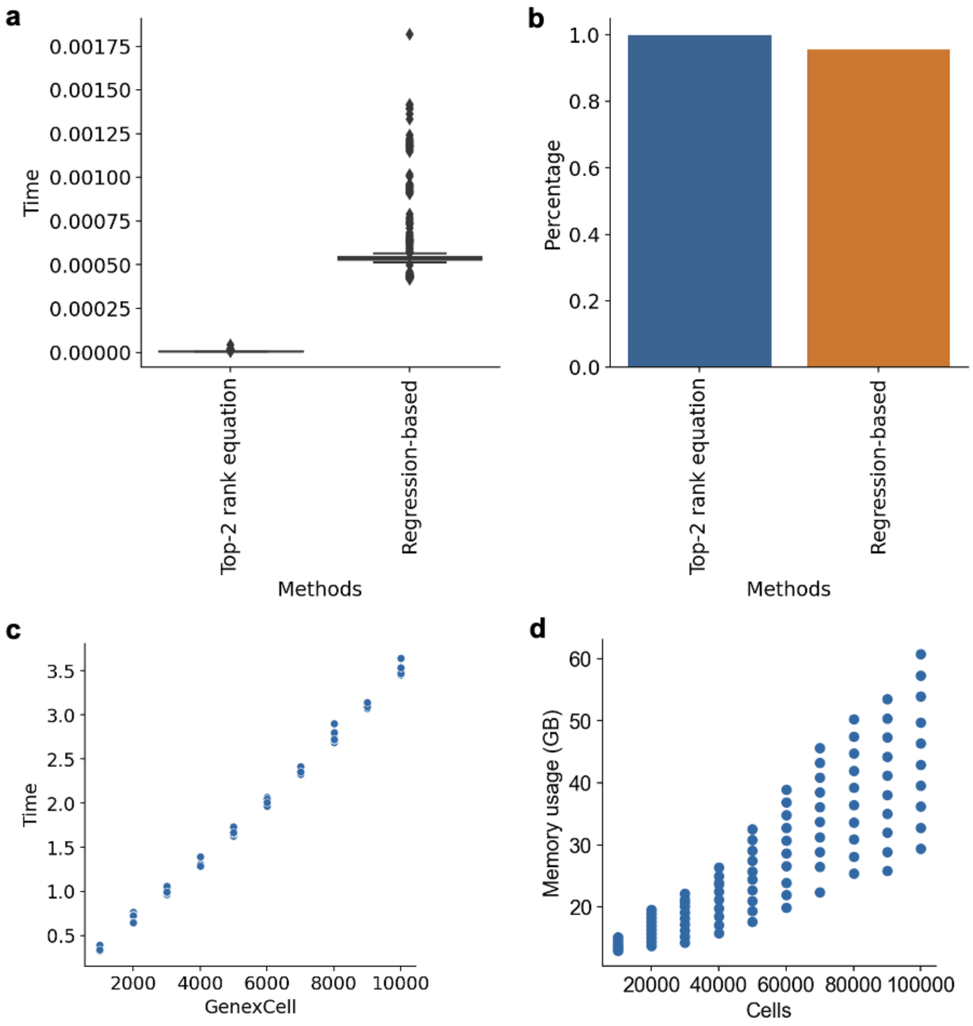


#### Supplementary Fig. 4 The computational efficiency of scDenorm

**a**, The boxplot shows the time distribution to calculate the scaling factors for cells by two unscaling methods, the top-2 rank equation method and the regression based method.

**b**, The histogram shows the percentage of cells that were successfully denormalised by the two methods.

**c**, The distribution of execution times for denormalisation (y-axis, the unit is second) across cells with genes ranging from 2000 to 10000 (x-axis).

**d**, The distribution of memory usage during denormalisation (y-axis, the unit is GB) across cells ranging from 10000 to 100,000 (x-axis).

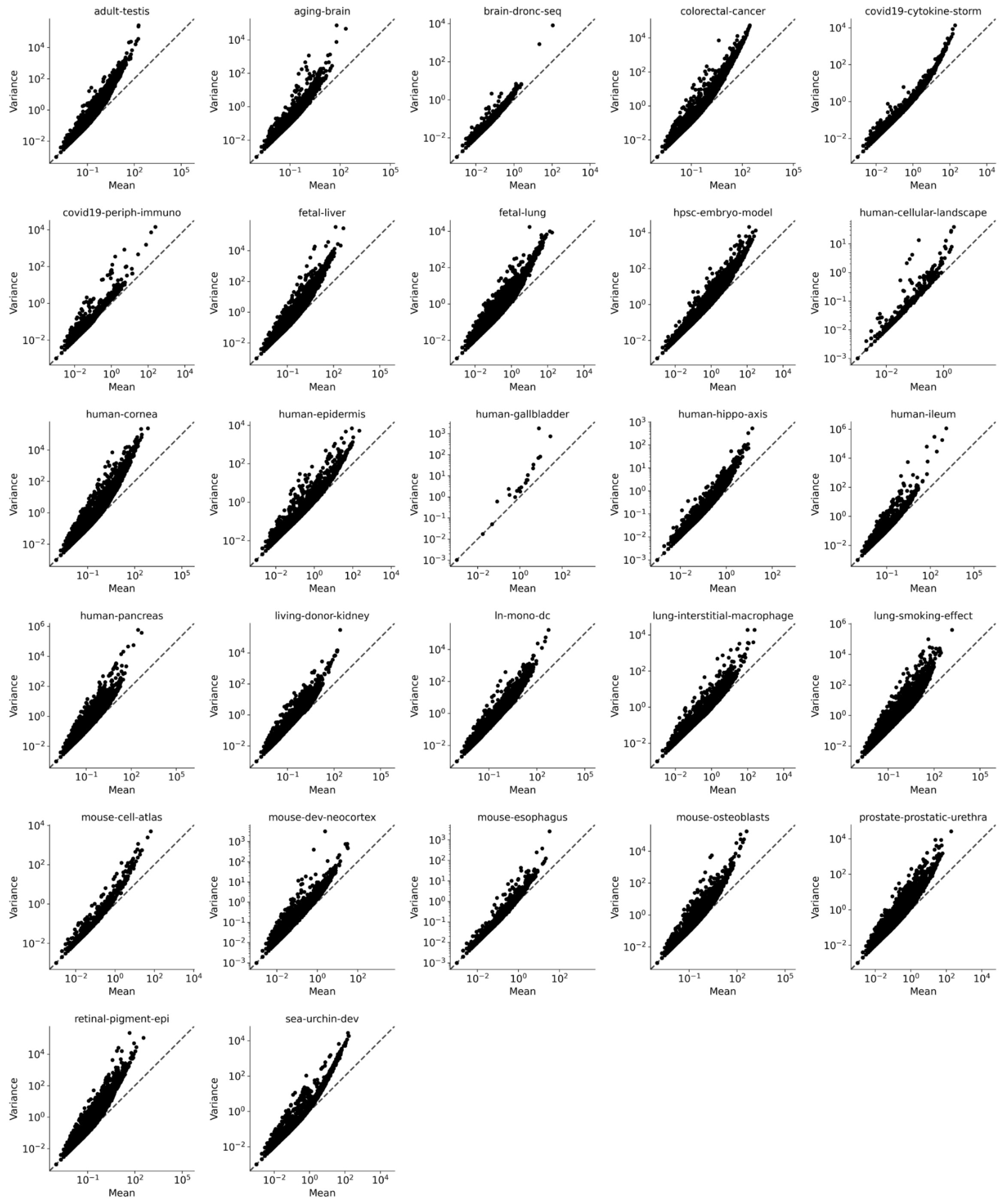


### Supplementary Fig. 5 The relationship between the mean and the variance of genes after denormalisation

Scatter plots show the relationship between log-transformed mean expression (x-axis) with log-transformed variance (y-axis) for each gene after denormalisation, across different scRNA-seq datasets from UCSC Cell Browser. Each dot represents a gene, with titles indicating the dataset names.

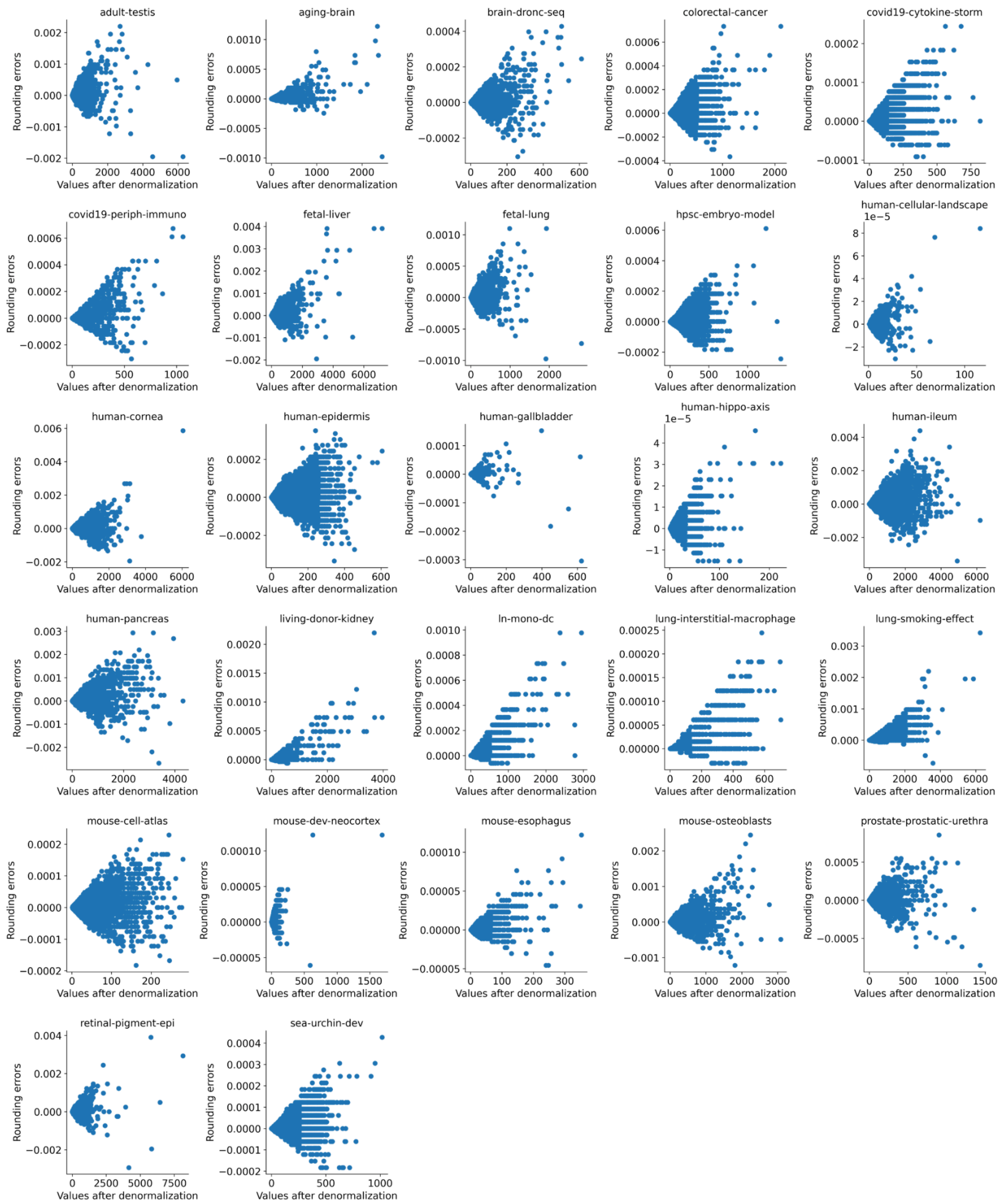


### Supplementary Fig. 6 The relationship between the rounding errors and values after denormalisation

Scatter plots illustrate the relationship between count values (x-axis) and rounding errors (y-axis) after denormalisation for different scRNA-seq datasets from UCSC Cell Browser. Each dot represents a cell, with titles indicating dataset names.

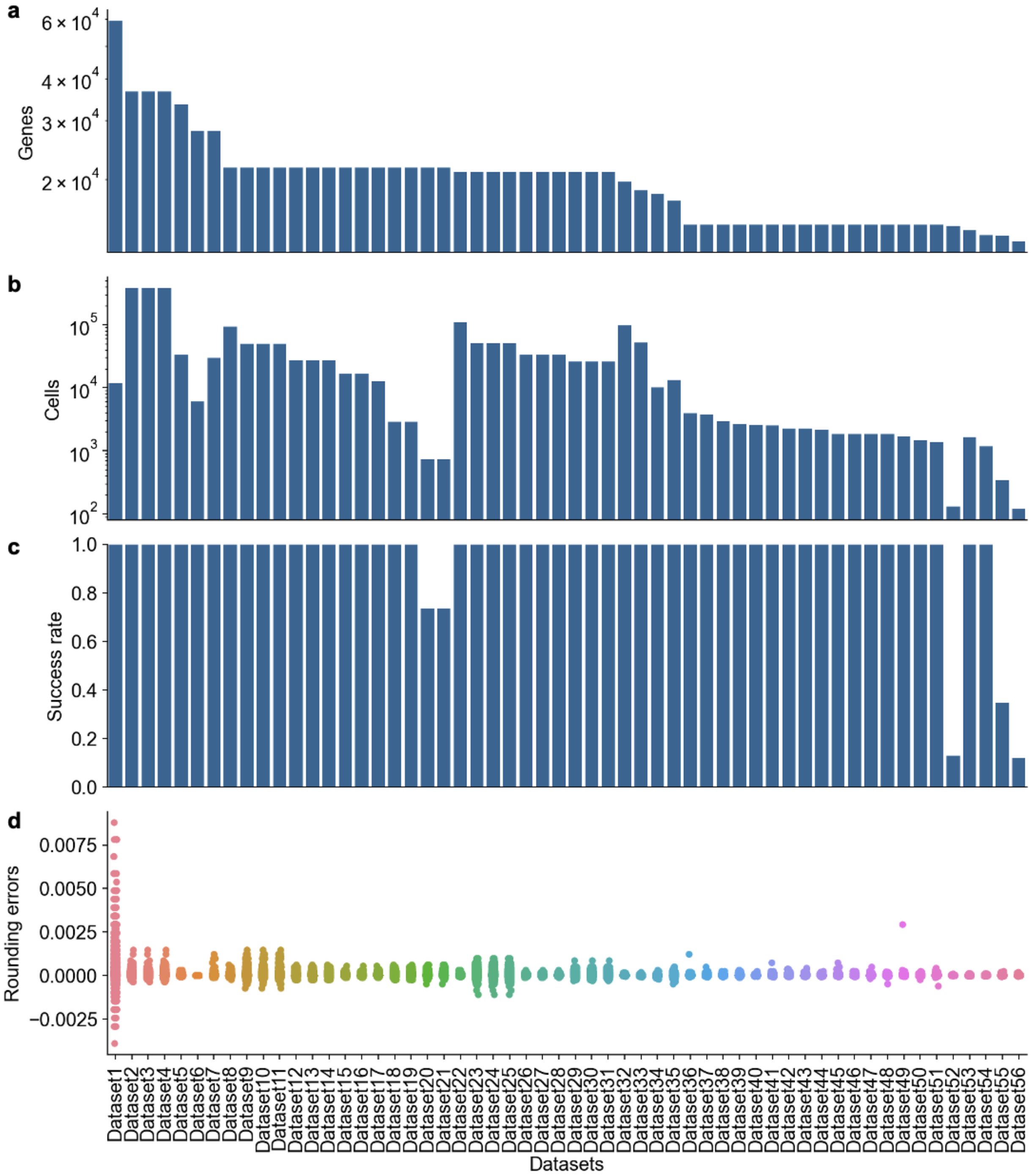


#### Supplementary Fig. 7 Benchmark analysis on the datasets from Brain Cell Atlas

**a,** The bar plot shows the number of genes in each dataset. The x-axis is the name of datasets (same as **d**), and the y-axis is the log-scaled number of genes.

**b,** The bar plot shows the number of cells in each dataset. The x-axis is the name of datasets

(same as **d**), and the y-axis is the log-scaled number of cells.
**c,** The bar plot shows the success rate for each dataset. The x-axis is the name of datasets (same as **d**), and the y-axis is the success rate (see **Method**).
**d,** The jitter plot shows the distribution of rounding errors observed in the denormalised datasets from the Brain Cell Atlas. The x-axis is the name of datasets, and the y-axis is the rounding error.

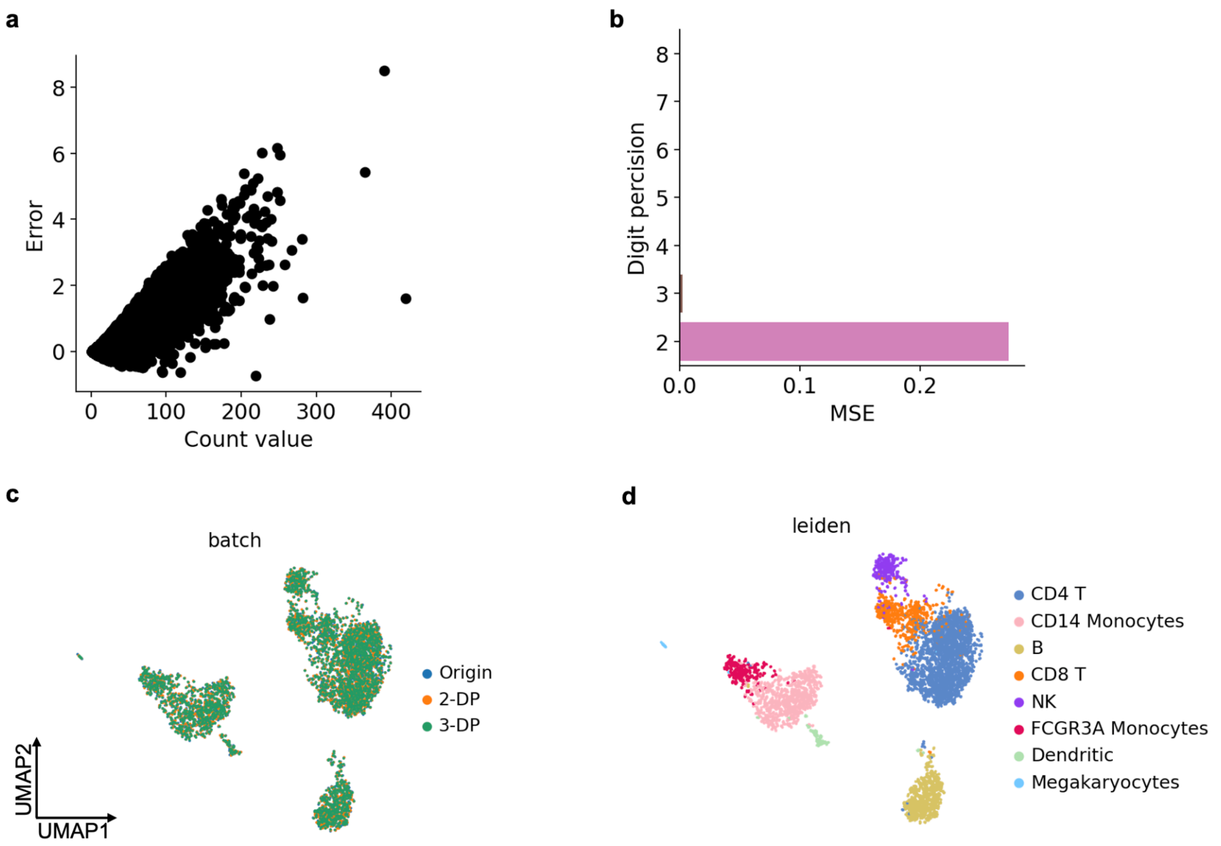


#### Supplementary Fig. 8 The robustness of scDenorm recovering raw counts in different scenarios

**a**, The relationship between the count values and the rounding errors after denormalisation on 2 digit precision data.
**b**, The mean square error of recovery error after denormalisation on the digit precisions from 2 to 8.
**c**, UMAP plot with different colors to represent different digit precisions, including original data, 2 digital precision data and 3 digital precision data.
**d**, UMAP plot (same as **c**) with different colors to represent different cell types.

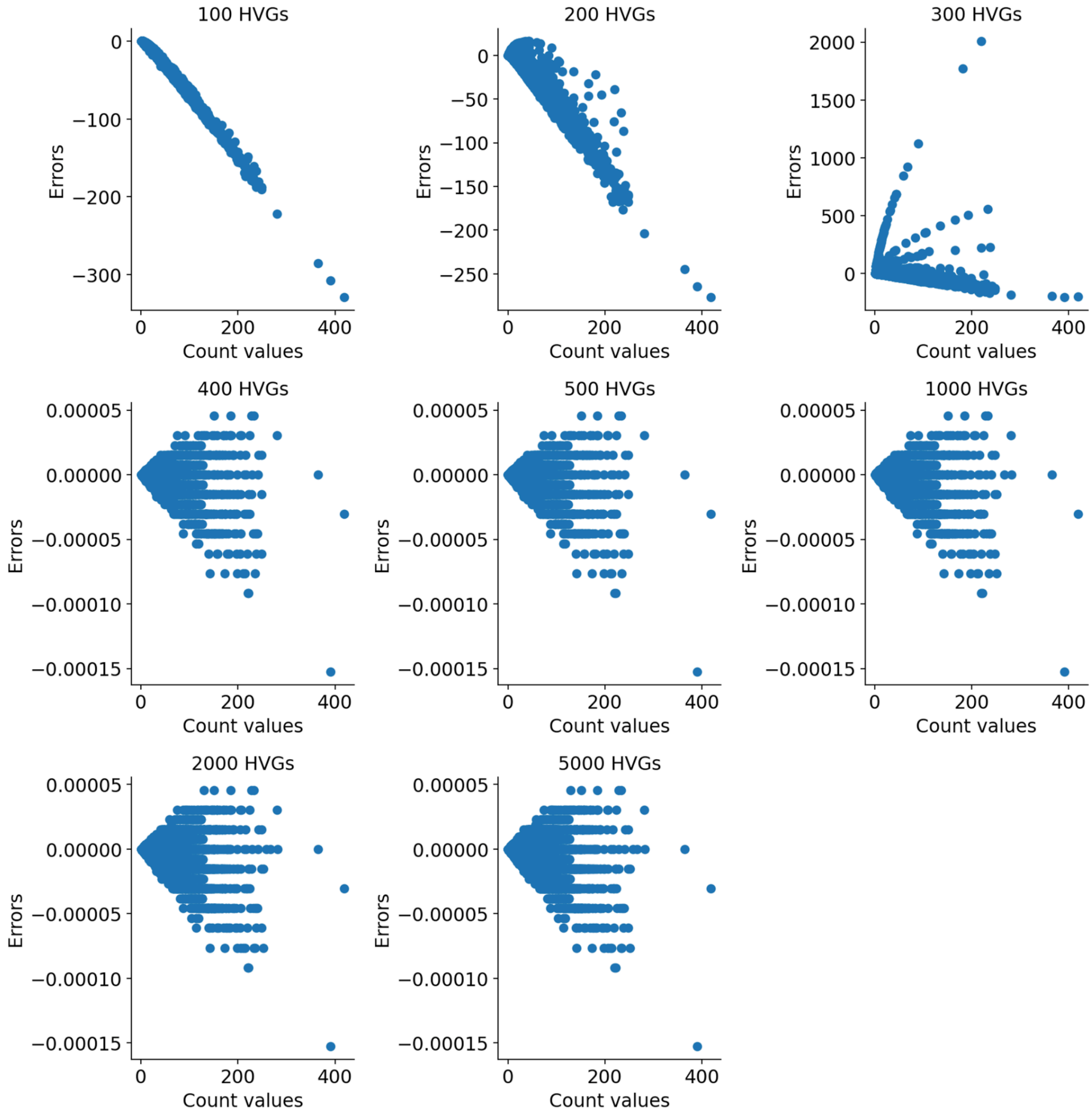


Supplementary Fig. 9 The relationship between the rounding errors and count values in different numbers of top highly variable genes

The scatter plots show that The relationship between count values (x-axis) and rounding errors (y-axis) after denormalisation, for different numbers of top highly variable genes. Each dot is a cell. The titles are the number of top highly variable genes.

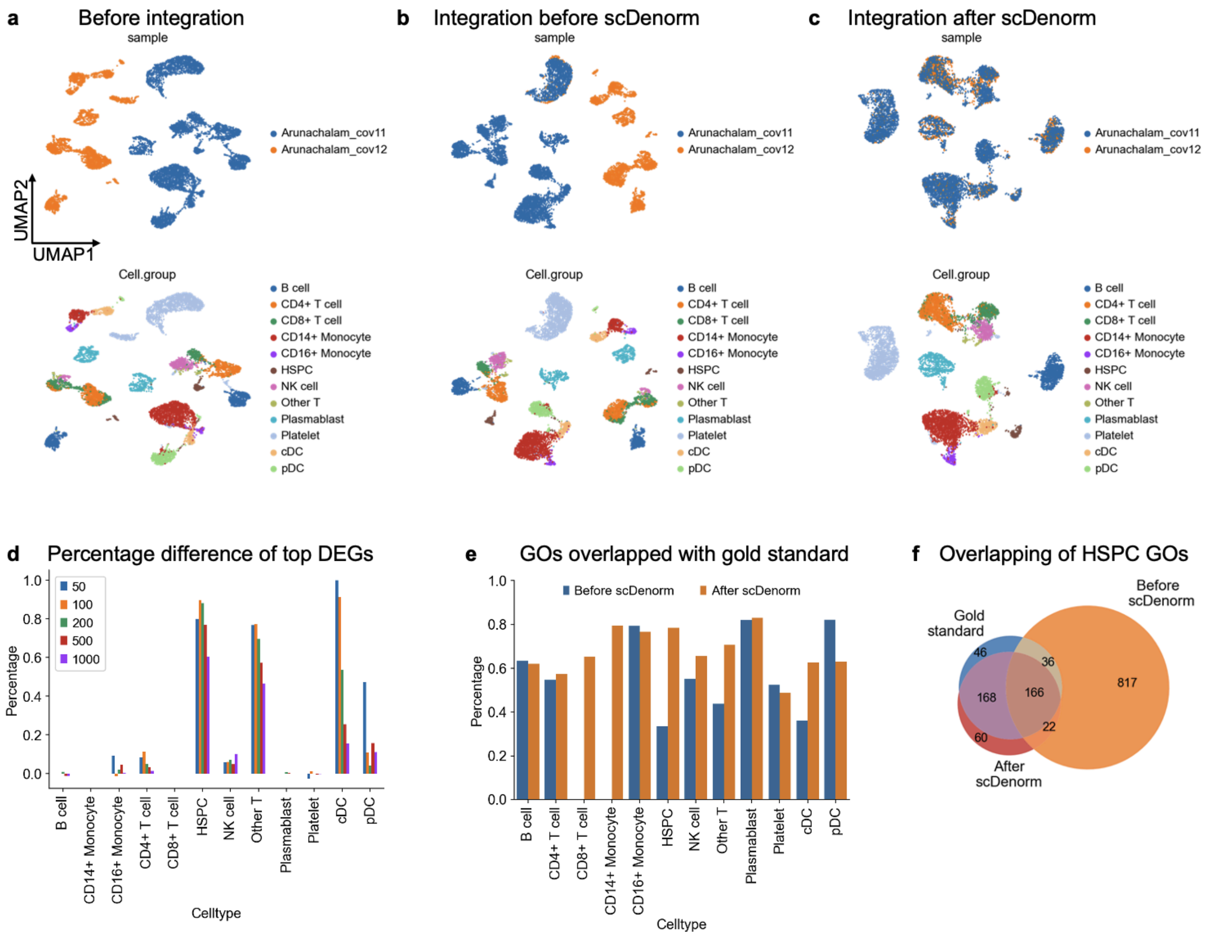


### Supplementary Fig. 10 The integration and downstream analyses on COVID-19 PBMC datasets

**a,** The UMAP plots show the distribution of cells of the COVID-19 PBMC datasets before integration without the scDenorm denormalisation, coloured by sample (top) , and original cell type from Arunachalam et al. (bottom).

**b,** The UMAP plots show the Harmony-integrated results without the scDenorm denormalisation, coloured by sample (top) , and original cell type from Arunachalam et al. (bottom).

**c,** The UMAP plots show the Harmony-integrated results after running scDenorm, coloured with sample (top) , and original cell type from Arunachalam et al. (bottom).

**d,** Bar plot shows the percentage difference of DEGs before and after scDenorm across cell types. The top DEGs were compared across different thresholds (top 50, 100, 200, 500, and 1000).

**e,** Bar plot shows the overlapping percentage of GO terms between the gold standard with before and after scDenorm across cell types.

**f,** Venn diagram shows the overlap of GO terms for HSPC among the gold standard, before scDenorm, and after scDenorm.

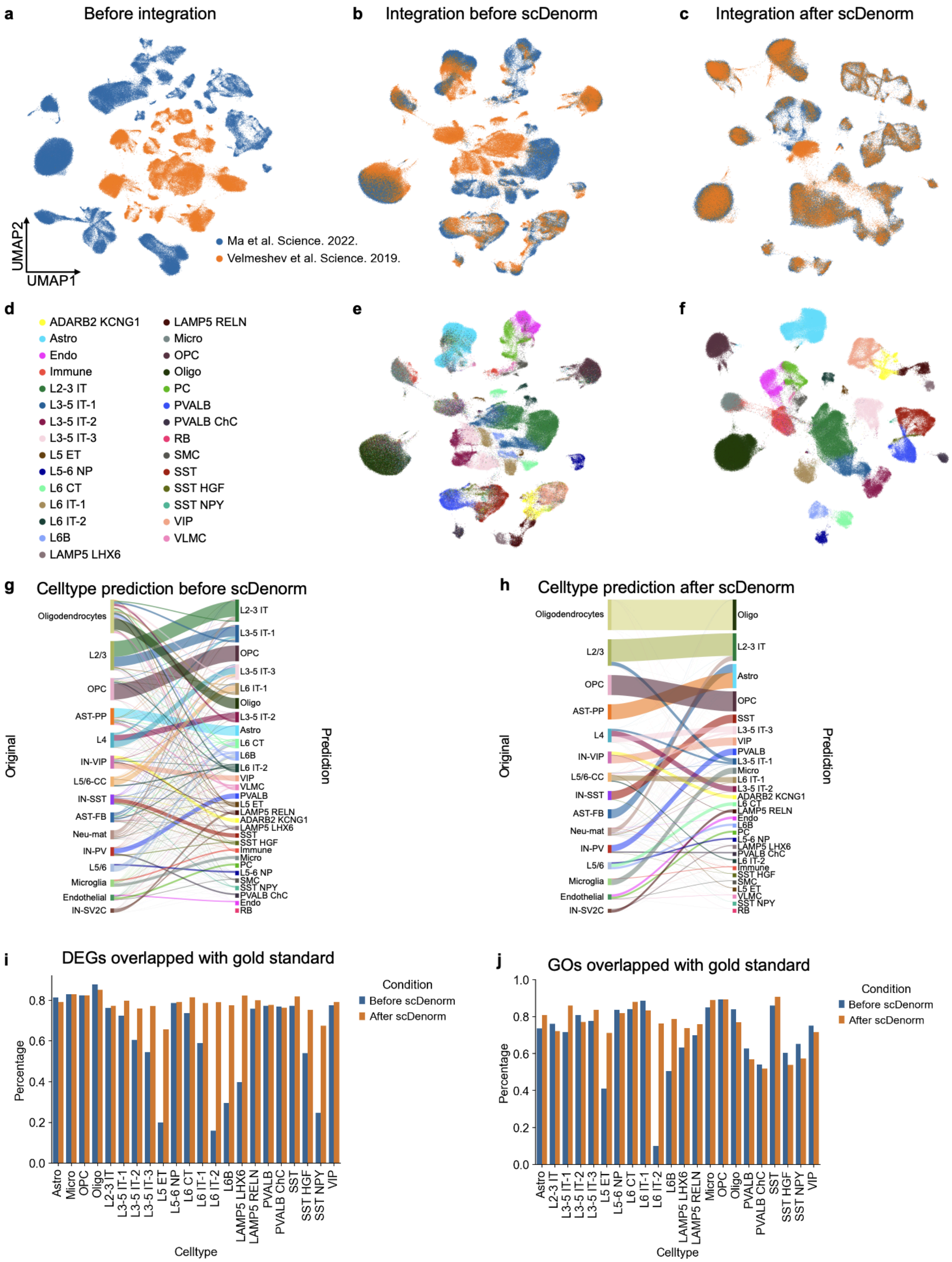
a
Before integration
b
Integration before scDenorm
c
Integration after scDenorm
UMAP2
UMAP1
Ma et al. Science. 2022.
Velmeshev et al. Science. 2019.
d
ADARB2 KCNG1
Astro
Endo
Immune
L2-3 IT
L3-5 IT-1
L3-5 IT-2
L3-5 IT-3
L5 ET
L5-6 NP
L6 CT
L6 IT-1
L6 IT-2
L6B
LAMP5 LHX6
LAMP5 RELN
Micro
OPC
Oligo
PC
PVALB
PVALB ChC
RB
SMC
SST
SST HGF
SST NPY
VIP
VLMC
e
f
g
Celltype prediction before scDenorm
Original
Prediction
Oligodendrocytes
L2/3
OPC
AST-PP
L4
IN-VIP
L5/6-CC
IN-SST
AST-FB
Neu-mat
IN-PV
L5/6
Microglia
Endothelial
IN-SV2C
L2-3 IT
L3-5 IT-1
OPC
L3-5 IT-3
L6 IT-1
Oligo
L3-5 IT-2
Astro
L6 CT
L6B
L6 IT-2
VIP
VLMC
PVALB
L5 ET
LAMP5 RELN
ADARB2 KCNG1
LAMP5 LHX6
SST
SST HGF
Immune
Micro
PC
L5-6 NP
SMC
SST NPY
PVALB ChC
Endo
RB
h
Celltype prediction after scDenorm
Original
Prediction
Oligodendrocytes
L2/3
OPC
AST-PP
L4
IN-VIP
L5/6-CC
IN-SST
AST-FB
Neu-mat
IN-PV
L5/6
Microglia
Endothelial
IN-SV2C
Oligo
L2-3 IT
Astro
OPC
SST
L3-5 IT-3
VIP
PVALB
L3-5 IT-1
Micro
L6 IT-1
L3-5 IT-2
ADARB2 KCNG1
L6 CT
LAMP5 RELN
Endo
L6B
PC
L5-6 NP
LAMP5 LHX6
PVALB ChC
L6 IT-2
Immune
SST HGF
SMC
L5 ET
VLMC
SST NPY
RB
i
DEGs overlapped with gold standard
Percentage
0.0
0.2
0.4
0.6
0.8
Condition
Before scDenorm
After scDenorm
Astro
Micro
OPC
Oligo
L2-3 IT
L3-5 IT-1
L3-5 IT-2
L3-5 IT-3
L5 ET
L5-6 NP
L6 CT
L6 IT-1
L6 IT-2
L6B
LAMP5 LHX6
LAMP5 RELN
PVALB
PVALB ChC
SST
SST HGF
SST NPY
VIP
Celltype
j
GOs overlapped with gold standard
Percentage
0.0
0.2
0.4
0.6
0.8
1.0
Condition
Before scDenorm
After scDenorm
Astro
L2-3 IT
L3-5 IT-1
L3-5 IT-2
L3-5 IT-3
L5 ET
L5-6 NP
L6 CT
L6 IT-1
L6 IT-2
L6B
LAMP5 LHX6
LAMP5 RELN
Micro
OPC
Oligo
PVALB
PVALB ChC
SST
SST HGF
SST NPY
VIP
Celltype

### Supplementary Fig. 11 scDenorm helps in cell type annotation and differential expression analysis on prefrontal cortex datasets

**a,** The UMAP plot shows the distribution of cells of the two prefrontal cortex datasets before integration.
**b,** The UMAP plot shows Harmony-integrated results without the scDenorm denormalisation, coloured by studies.
**c,** The UMAP plot shows the Harmony-integrated result after running scDenorm, coloured by studies.
**d,** The figure legend of cell type annotation for panel (**e**) and (**f**).
**e,** The UMAP plot is the same as (**b**) coloured by CellTypist-predicted cell type annotation.
**f,** The UMAP plot is the same as (**c**) coloured by CellTypist-predicted cell type annotation.
**g-h,** River plot illustrates the transition between original and predicted cell types before scDenorm (**g**) and after scDenorm (**h**). The left side represents the original cell types from Velmeshev et al., while the right side displays the predicted cell types.
**i,** Bar plot showing the overlapping percentage of DEGs between the gold standard with before and after scDenorm across cell types. The DEGs are calculated with a two-sided Wilcoxon test based on the predicted cell types.
**j,** Bar plot showing the overlapping percentage of GO terms between the gold standard with before and after scDenorm across cell types.

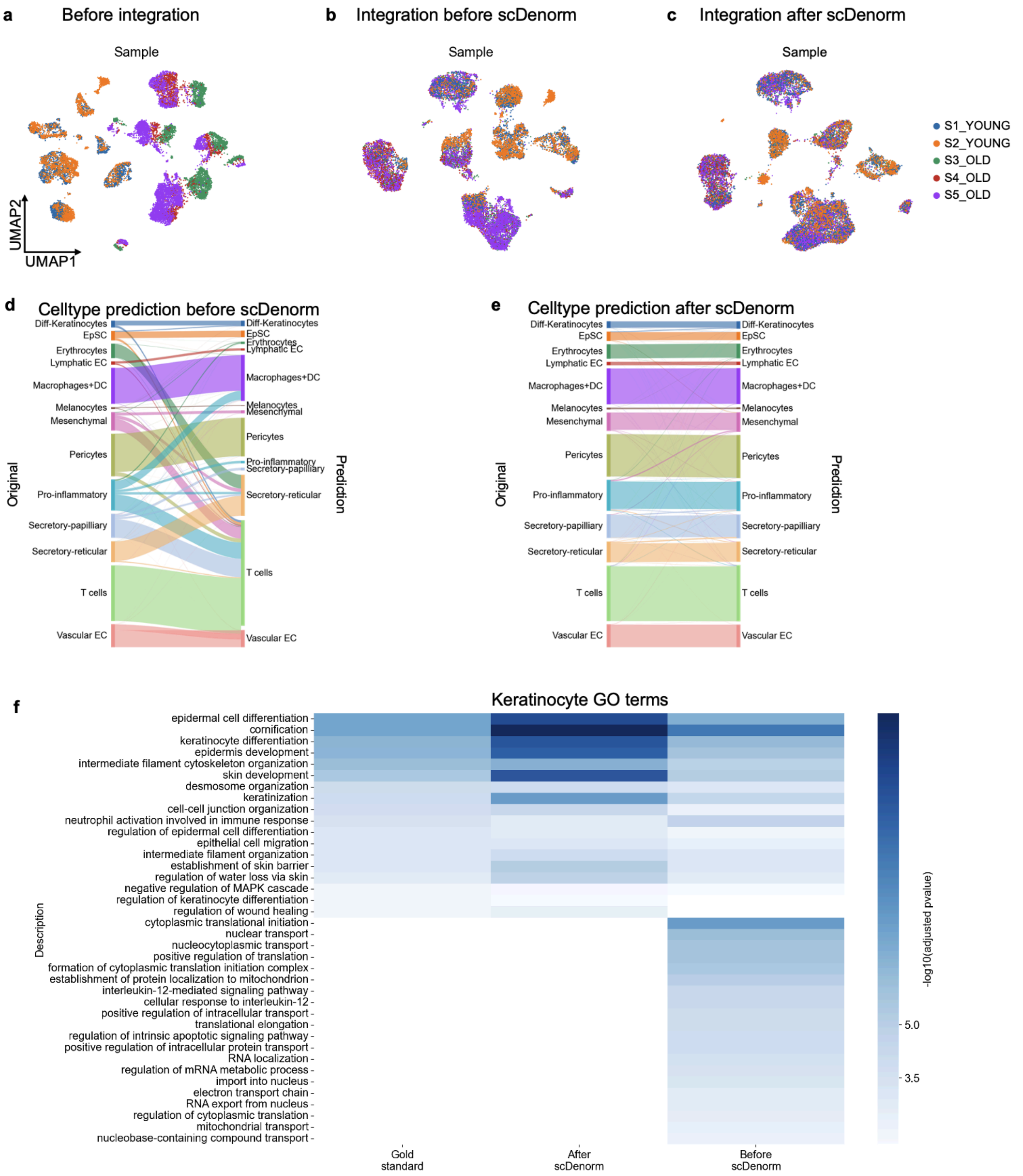


### Supplementary Fig. 12 scDenorm facilitates the downstream analyses on young and old human skin datasets

**a,** The UMAP plot shows the cell distribution of the human skin datasets before integration without the scDenorm denormalisation, coloured with samples.

**b,** The UMAP plot shows the Harmony-integrated result without the scDenorm denormalisation,

coloured by samples.
**c,** The UMAP plot shows the Harmony-integrated result after running scDenorm, coloured by samples.
**d,** River plot illustrates the transition between original and predicted cell types before scDenrom. The left side represents the original cell types from Solé-Boldo et al., while the right side displays the predicted cell types.
**e,** River plot illustrates the transition between original and predicted cell types after scDenrom. The left side represents the original cell types from Solé-Boldo et al., while the right side displays the predicted cell types.
**f,** Heatmap shows the enriched Gene Ontology (GO) terms for the keratinocyte's DEGs identified in the gold standard, before scDenorm as well as after scDenorm.

### Supplementary Tables

#### Supplementary Table 1. Description of the datasets from UCSC Cell Browser.

#### Supplementary Table 2. Description of the datasets from Brain Cell Atlas.

#### Supplementary Table 3. Summary of the normalisation methods on 133 well-established single-cell studies.
