## Supplementary material for "scDenorm: a denormalisation tool for integrating single-cell transcriptomics data": Rebuttal

Dear Veronique,

We gratefully appreciate your feedback and all the comments from the reviewers. The constructive feedback helped us to improve our manuscript. We have carefully addressed the concerns raised by the reviewers and have made substantial improvements to the manuscript. Please find our point-by-point responses below:

| **Comments** | **Answers** |
|---|---|
| Editor | |
| **0.0** Thank you very much for submitting your manuscript entitled 'scDenorm: a denormalisation tool for Integrating Single-cell Transcriptomics Data' to Genome Biology, and please accept my apologies for the delay in replying to you about it. It has now been seen by two referees and their comments are accessible below. Please also take a moment to check our website at https://www.editorialmanager.com/gbio/ for additional comments by Referee #2 saved as attachment.<br>As you will see from the reports, the referees find the manuscript of potential interest, but they raise serious concerns that you have not convinced about the utility of scDenorm. In particular, for further consideration at Genome Biology you | Thank you for the feedback! We have now systematically revised our manuscript according to the suggestions from the reviewers. More complete benchmark studies are provided for validation. Please find all these improvements below in our point-to-point answers to the reviewers' comments. |

| | |
|---|---|
| should absolutely convince of the necessity and advance of applying scDenorm as suggested by both referees. In addition to demonstrating biological insights are missed by not applying scDenorm, you should expand the validation and also the benchmarking by adding more datasets and metrics. It seems to us to be essential that all of the referees' concerns are fully addressed, in the form of a revised manuscript, before we can reach a final decision on publication.<br><br>We will only send the manuscript back out for re-review if we feel that the concerns highlighted above have been sufficiently addressed. | |
| **0.1** We are considering your manuscript as a 'Method' article. This is our format for publishing articles that describe a methodological innovation that is a significant advance over published methods and likely to be of broad utility, but that do not provide significant biological insights. When revising the manuscript, please ensure the manuscript conforms to our style for Methods articles (see | Thanks. We have ensured that the manuscript conforms to your style guidelines, including the 100-word limit for the abstract, adding the conclusion section. The source codes are now publicly available under an OSI-compliant open source license as required. |

| | |
|---|---|
| https://genomebiology.biomedcentral.com/submission-guidelines/preparing-your-manuscript/method); specifically, the abstract should be under 100 words. Please note that if we decide to publish your manuscript we will require that the source code is made publicly available under an open source license compliant with Open Source Initiative, with the license clearly stated in the manuscript. | |
| **0.2** The source code should be deposited in a public repository, such as for instance github, with the accession links included in the manuscript. We also ask that the version of source code used in the manuscript is deposited in a DOI-assigning repository, such as zenodo, with the link also included. All this information should be listed in a separate Availability of Data and Materials section of the manuscript. | The source codes of the scDenorm program are publicly available at GitHub repo (scDenorm):<br>https://github.com/rnacentre/scDenorm<br><br>And codes to reproducing the figures in this manuscript are available at GitHub repo (scDenorm-reporoducibility):<br>https://github.com/rnacentre/scDenorm-reproducibility<br><br>Additionally, the dataset related to this manuscript can be accessed via Zenodo: https://doi.org/10.5281/zenodo.11240443 |
| Reviewer 1 | |
| **1.0**<br>Normalization plays a pivotal role in precisely comparing gene expression | Thanks! Following the reviewers' suggestions, we have now made significant revisions of our manuscript. The highlighted changes are listed: |

| | |
|---|---|
| levels among cells or samples. However, given the diverse normalization approaches adopted for different datasets, direct integration of data may inadvertently introduce errors. The author team developed scDenorm to address this issue by converting normalized expression data to row counts. The idea of de-normalization is innovative and this algorithm is relatively concise and should be quite effective. However, this algorithm is limited to de-normalize data that normalized by delta method, thereby restricting its potential applications. In addition, I think the importance of this work should be further emphasized and figures need substantial improvement before I can recommend the publication of this manuscript. | 1. More complete benchmarks and downstream analysis validation.<br>In the revised version, we provide further validations of our scDenorm approach, including: **a**) A survey of well-established single-cell studies reports that 83% of the datasets are normalised with delta method, indicating the application scope of scDenorm; **b**) Further large-scale benchmark test on 60 human brain datasets suggests that 93% of the datasets can be denormalised with scDenorm; **c**) The denormalisation effects of scDenorm was benchmarked on downstream analysis tasks, including data integration, cell type annotation, differential gene expression and gene ontology analysis. The three benchmark datasets cover different data batch types, including sample batch, study batch and condition batch; **d**) The computational time and memory consumptions were benchmarked. In addition, more quantifiable metrics are used for all these above mentioned benchmarks.<br><br>2. The clarity of the methodology was improved.<br>In the Methods section, we a) improved the definition of all the terms and equations; b) included more details about evaluation metrics, data processing and parameter optimisation.<br><br>3. The manuscript together with all the figures were systematically revised.<br>We systematically revised the manuscript, included more results descriptions and discussions, and optimised all the figures for clarity. |
| **1.1** Currently, the manuscript merely demonstrates that de-normalization has an impact on data integration, but I believe that this alone is insufficient to convey the significance of this work. Therefore, I propose investigating whether the application of scDenorm not only impacts data integration but also downstream biological | Thanks for the nice suggestion. Three additional case studies were included to highlight the normalisation-resulted biases in downstream analysis, including data integration, cell type annotation, differential expression and gene ontology analysis. These three datasets cover different batch types, including sample batch, study batch and condition batch. The downstream analysis steps are discussed as below:<br><br>1. Misassignment in cell type annotation:<br>Reference-based cell type annotation based on machine learning is a well-established approach, e.g., CellTypist. It assigns the cell types according to the comparison between |

| | |
|---|---|
| discoveries. Is there example to illustrate how not using scDenorm may lead to incorrect biological conclusions? | the expression profiles in the reference dataset and the target dataset. We show that different normalisations may result in differences in the expression profile, thus leading to mis-annotated cell types. CellTypist, which is a logistic regression based approach, was used to annotate cell types before and after scDenorm on three datasets, including COVID-19 PBMC dataset (see **Fig. 5**), human skin dataset (see **Fig. 6**), and human prefrontal cortex dataset (see **Supplementary Fig. 11**). The CellTypist-assigned cell type annotation after scDenorm shows better consistency with the cell type label reported in the original publication than CellTypist-assigned annotation according to normalised data before scDenorm. The annotation accuracy increases from 66% to 92% (after scDenorm). And such mislabeled cell types may also lead to misinterpretation in downstream analysis, such as differential expression analysis and enrichment analysis. See P6L222.<br><br>2. Biases in Differential Expression (DE) analysis:<br>Taking the published differential expression genes as ‘gold standard’, we performed one-vs.-rest DE analysis for each cell type to derive the cell type specific DEs before and after scDenorm. We conducted DE analysis between relevant cell types on the human skin dataset (see **Fig. 6**), comparing the results obtained before and after scDenorm. DE analysis after scDenorm domenstrates higher consistency with the 'gold standard' (**Fig. 6h**), and fewer false-positive differentially expressed genes (**Fig. 6i**).<br><br>3. Misinterpretations in functional enrichment and biological relevance:<br>Building on the DE analysis, we performed Gene Ontology analysis to understand the biological implications. We compared the enriched pathways, and processes when using the dataset before and after scDenorm. GO analysis shows that the GO terms associated with data before scDenorm have low consistency with 'gold standard' terms derived from the ‘gold standard’ DEGs from the original publication (**Fig. 6h**). For instance, GO analysis of the differential expression genes in HSPCs show that the GO terms enriched after scDenorm are well aligned with those from the 'gold standard', whereas the enriched terms before scDenorm showed minimal overlap with 'gold standard' (**Supplementary Fig. 10f**). The enriched GO terms after scDenorm are |

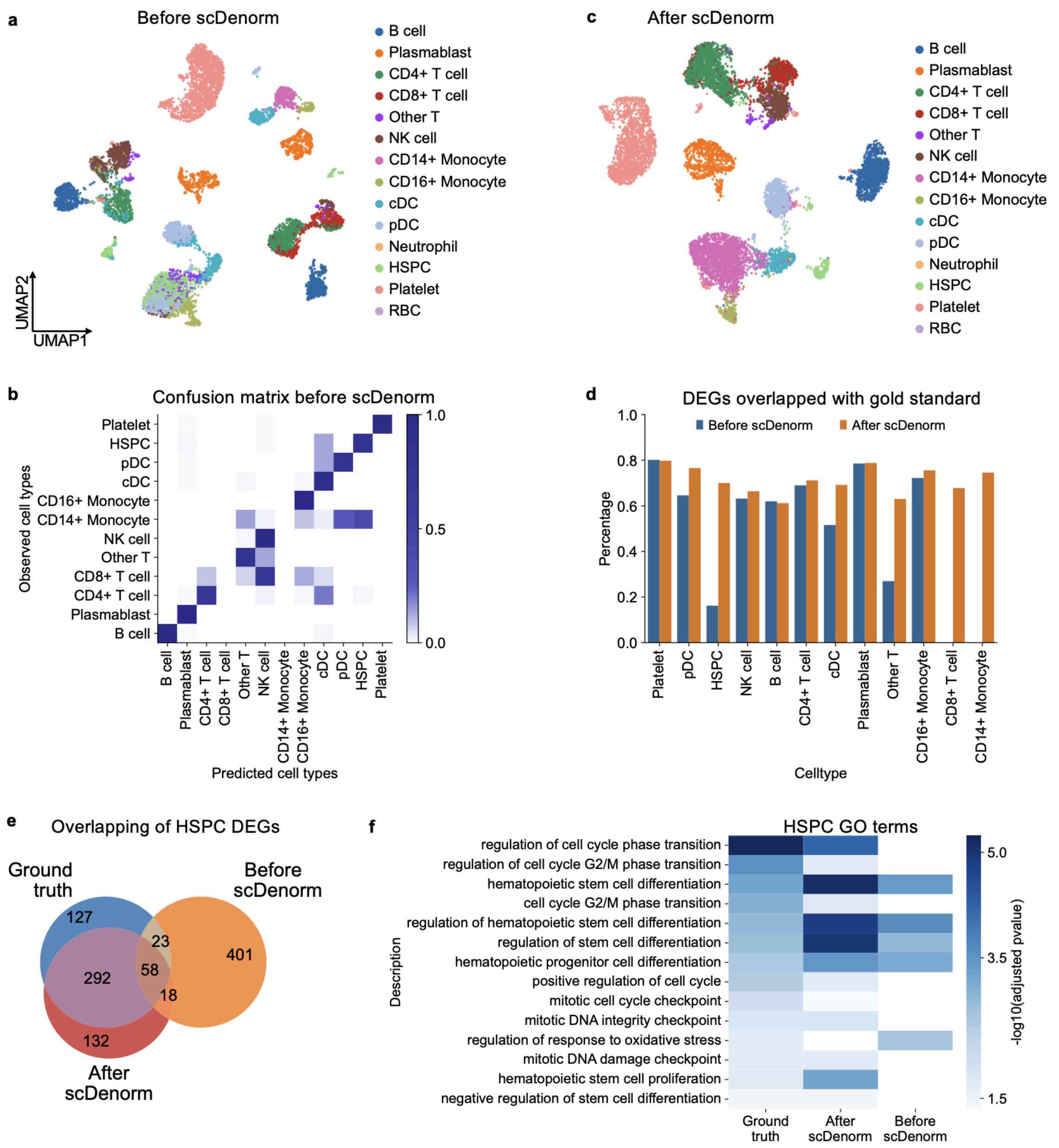
a
Before scDenorm
B cell
Plasmablast
CD4+ T cell
CD8+ T cell
Other T
NK cell
CD14+ Monocyte
CD16+ Monocyte
cDC
pDC
Neutrophil
HSPC
Platelet
RBC
UMAP2
UMAP1
c
After scDenorm
b
Confusion matrix before scDenorm
Observed cell types
Predicted cell types
d
DEGs overlapped with gold standard
Before scDenorm
After scDenorm
Percentage
Celltype
e
Overlapping of HSPC DEGs
Ground truth
Before scDenorm
After scDenorm
127
23
401
292
58
18
132
f
HSPC GO terms
regulation of cell cycle phase transition
regulation of cell cycle G2/M phase transition
hematopoietic stem cell differentiation
cell cycle G2/M phase transition
regulation of hematopoietic stem cell differentiation
regulation of stem cell differentiation
hematopoietic progenitor cell differentiation
positive regulation of cell cycle
mitotic cell cycle checkpoint
mitotic DNA integrity checkpoint
regulation of response to oxidative stress
mitotic DNA damage checkpoint
hematopoietic stem cell proliferation
negative regulation of stem cell differentiation
Description
Ground truth
After scDenorm
Before scDenorm
-log10(adjusted pvalue)

| | |
|---|---|
| | **Fig. 5 Different normalisations impact cell type annotation of COVID-19 PBMCs**<br>**a,** The UMAP plot shows the distribution of cells before denormalisation by scDenorm, coloured by CellTypist-predicted cell type annotation (see **Methods**).<br>**b,** The UMAP plot shows the distribution of cells after Harmony data integration following by scDenorm. Cells are coloured by CellTypist-predicted cell type annotation.<br>**c**. The heatmap shows the confusion matrix between the published cell type labels and the CellTypist-predicted cell types based on the Harmony-integrated latent space before scDenorm denormalisation. The x-axis represents predicted cell types, while the y-axis denotes original cell type annotation published in the study.<br>**d**. The histogram shows the percentage of DEGs overlap between the 'gold standard' (DEGs derived according to the original published cell type labels) and the ones derived from re-analysis before (blue) and after (orange) scDenorm across cell types. The DEGs are calculated with a two-sided Wilcoxon test using the CellTypist-predicted cell types as clusters.<br>**e**. The Venn diagram shows the overlap of the top 500 DEGs for HSPCs derived from the 'gold standard' (blue) from original study, as well as before (orange) and after (red) scDenorm.<br>**f**. The heatmap shows the enriched Gene Ontology (GO) terms of HSPCs derived from DEGs of the 'gold standard' (blue) from original study, as well as before (orange) and after (red) scDenorm. |

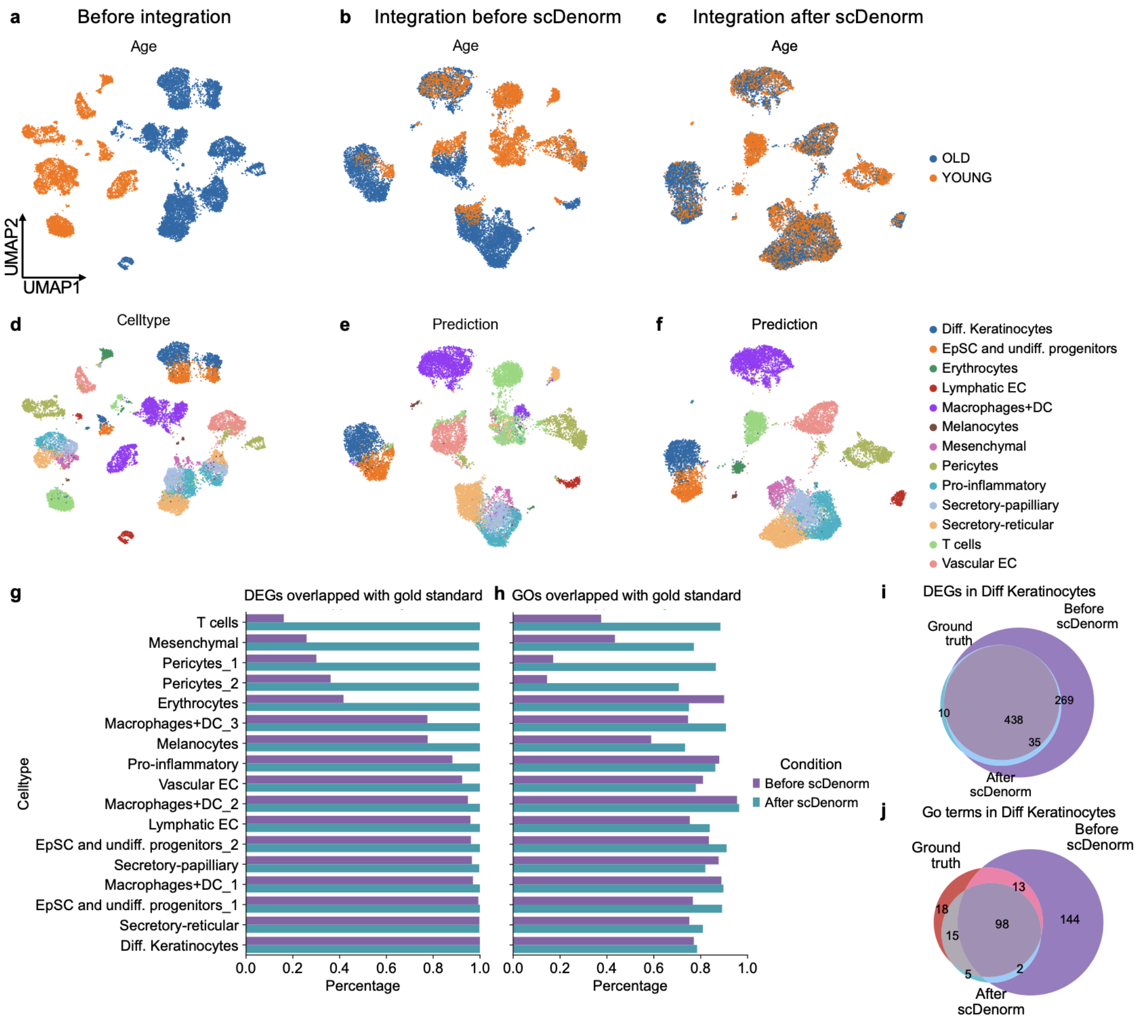


**Fig. 6 Different normalisations impact differential expression analysis**

**a,** The UMAP plot shows the cell distribution of the two age groups of the human skin

| | |
|---|---|
| | dataset (Solé-Boldo et al.) before data integration without the scDenorm denormalisation, coloured by age groups.<br>**b,** The UMAP plot shows the Harmony-integrated result without the scDenorm denormalisation, coloured by age groups.<br>**c,** The UMAP plot shows the Harmony-integrated result after running scDenorm, coloured by age groups.<br>**d,** The UMAP plots show the same distribution as panels (**a**), coloured by original cell type labels from Solé-Boldo et al.<br>**e-f,** The UMAP plots show the same distribution as panels (**b-c**), but coloured by CellTypist-predicted cell type labels before and after denormalisation by scDenorm.<br>**g-h,** The histograms show the percentages of DEGs (**g**) and GO terms (**h**) overlap between the 'gold standard' (DEGs extracted from the data in the original study, GO terms derived from these DEGs) and data before (purple) and after (blue) scDenorm across cell types, using the 'gold standard' cell type labels reported in the original publication. The DEGs are calculated with a two-sided Wilcoxon test based on the original cell type from the human skin dataset (Solé-Boldo et al.), while the GO analysis shows Benjamini-Hochberg-adjusted P value $<0.05$.<br>**i-j,** The Venn diagrams show the overlaps of the DEGs (**i**) (Benjamini-Hochberg-adjusted P value $<0.05$ and logFC $>0.25$) and GO terms (**j**) for differentiated keratinocytes derived from data of the 'gold standard' (red), as well as before (purle) and after (blue) scDenorm denormalisation. |

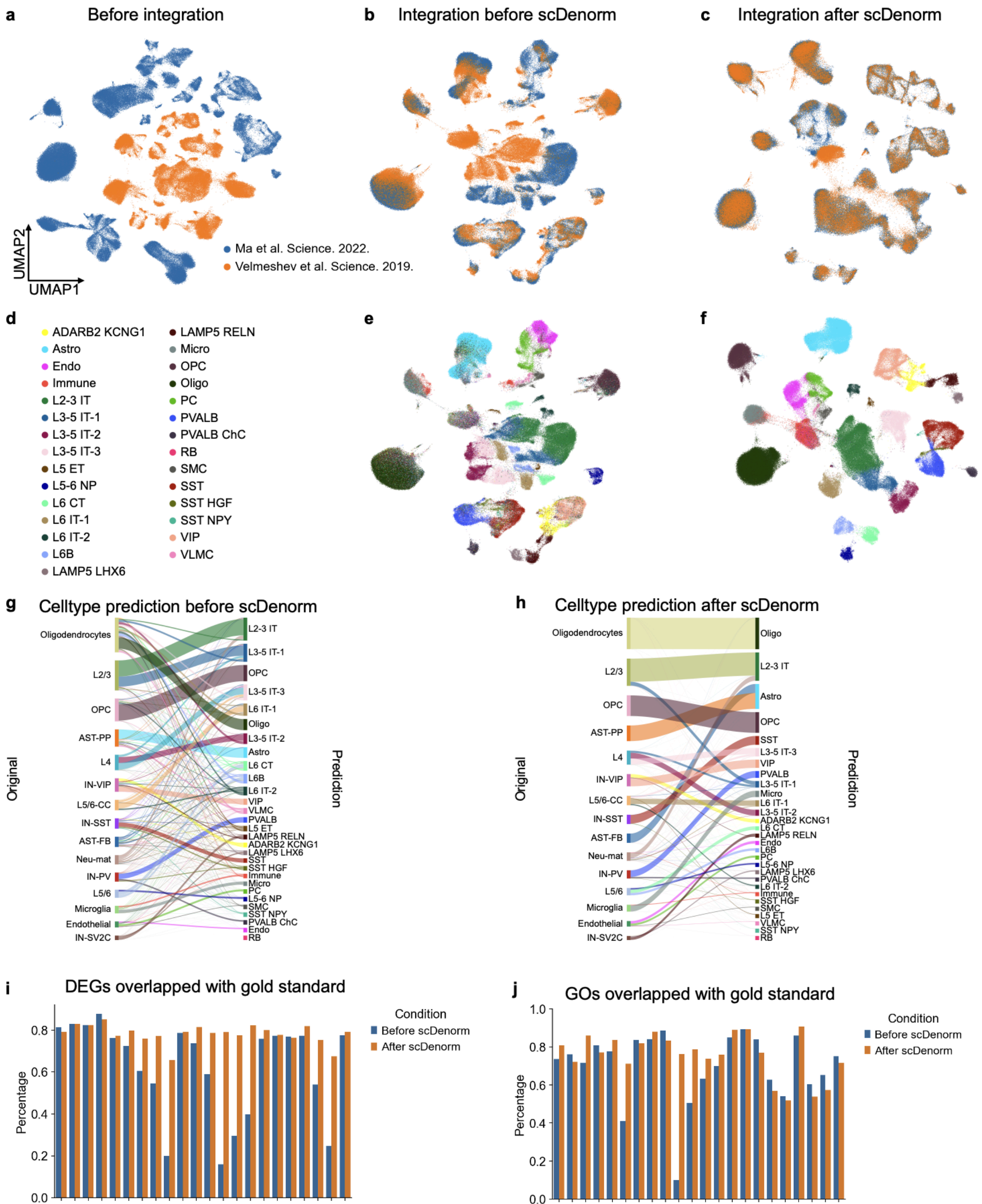

a
Before integration
b
Integration before scDenorm
c
Integration after scDenorm
UMAP2
UMAP1
Ma et al. Science. 2022.
Velmeshev et al. Science. 2019.
d
ADARB2 KCNG1
Astro
Endo
Immune
L2-3 IT
L3-5 IT-1
L3-5 IT-2
L3-5 IT-3
L5 ET
L5-6 NP
L6 CT
L6 IT-1
L6 IT-2
L6B
LAMP5 LHX6
LAMP5 RELN
Micro
OPC
Oligo
PC
PVALB
PVALB ChC
RB
SMC
SST
SST HGF
SST NPY
VIP
VLMC
e
f
g
Celltype prediction before scDenorm
Original
Prediction
h
Celltype prediction after scDenorm
i
DEGs overlapped with gold standard
Percentage
Condition
Before scDenorm
After scDenorm
j
GOs overlapped with gold standard

| | |
|---|---|
| | **Supplementary Fig. 11 scDenorm helps in cell type annotation and differential expression analysis on prefrontal cortex datasets**<br>**a,** The UMAP plot shows the distribution of cells of the two prefrontal cortex datasets before integration.<br>**b,** The UMAP plot shows Harmony-integrated results without the scDenorm denormalisation, coloured by studies.<br>**c,** The UMAP plot shows the Harmony-integrated result after running scDenorm, coloured by studies.<br>**d,** The figure legend of cell type annotation for panel (**e**) and (**f**).<br>**e,** The UMAP plot is the same as (**b**) coloured by CellTypist-predicted cell type annotation.<br>**f,** The UMAP plot is the same as (**c**) coloured by CellTypist-predicted cell type annotation.<br>**g-h,** River plot illustrates the transition between original and predicted cell types before scDenorm (**g**) and after scDenorm (**h**). The left side represents the original cell types from Velmeshev et al., while the right side displays the predicted cell types.<br>**i,** Bar plot showing the overlapping percentage of DEGs between the gold standard with before and after scDenorm across cell types. The DEGs are calculated with a two-sided Wilcoxon test based on the predicted cell types.<br>**j,** Bar plot showing the overlapping percentage of GO terms between the gold standard with before and after scDenorm across cell types. |
| **1.2** For de-transformation, the author presents two approaches, include (1) apply empirical values and (2) determine the parameters by solving optimization methods. For cells that meet the criteria in step (c), if scDenorm perform both (1) and (2), how consistent are the results | Thanks for the suggestion!<br><br>We benchmarked these two approaches (applying empirical values and solving optimization) on three datasets and found that the results are nearly identical, with discrepancies less than 0.01 (**Revision Fig. 1**). Trying empirical values is an efficient approach if empirical values were used in normalisation, while the solving optimization method (regression) is applicable to any parameters for delta method normalisation. |

<table>
<tr>
<td>obtained using the two methods? In addition, please show the detail methods you applied to solve the optimization problem.</td>
<td>
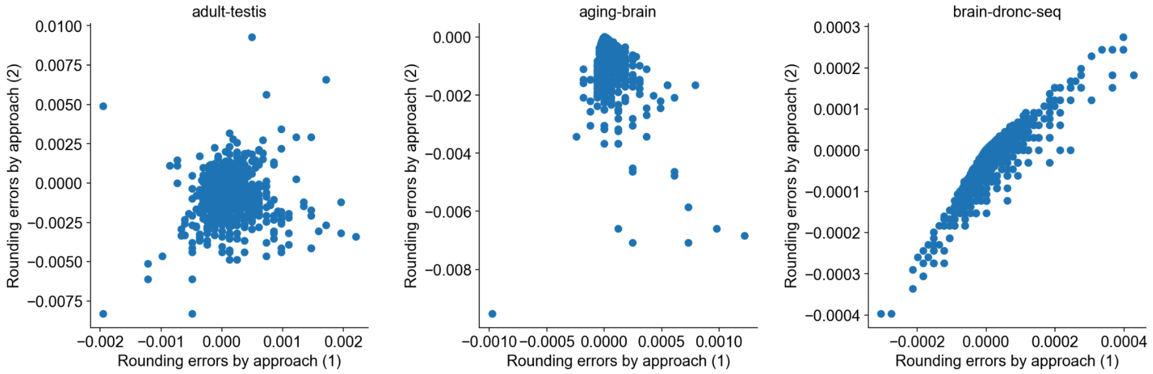


**Revision Fig, 1** The scatter plots between rounding errors from approach (1) apply empirical values and rounding errors from approach (2) determine the parameters by solving optimization methods.<br><br>We have now clarified in the Methods section about the optimisation method. Specifically, we utilized the L-BFGS-B method from the sklearn package to find the logarithmic base and pseudo count. This method is based on the limited-memory Broyden-Fletcher-Goldfarb-Shanno (BFGS) algorithm, which is particularly well-suited for large-scale optimization. L-BFGS-B allows for box constraints, which helps in ensuring that the parameters stay within specified bounds during optimization. See P10L387:<br><br>The L-BFGS-B method from the sklearn[43] package is used to find the best base (b) and pseudo count (p). This method is based on the limited-memory Broyden-Fletcher-Goldfarb-Shanno (BFGS) algorithm, which is capable of large-scale optimization. L-BFGS-B allows for box constraints, ensuring that the parameters stay within specified bounds during optimization.
</td>
</tr>
<tr>
<td>**1.3** If a single input h5ad file contains cells normalized with different empirical bases and pseudo counts,</td>
<td>Thanks for the suggestion! We have now implemented an iterative function ‘scdenorm’ with a ‘by=sample’ parameter as input to solve this problem. Considering that each sample in the same h5ad is normalised in the same set of parameters, we separate the</td>
</tr>
</table>

| | |
|---|---|
| will scDenorm work in this situation? | data by sample ID and use scDenorm to solve the parameters for each sample. After denormalising all the samples, we concatenate all the raw counts together. See P11L433:<br><br>Considering that different samples in a dataset may be normalised with different parameter sets, scDenorm also implements a per sample denormalisation function overloading the original ‘scdenorm’ function with a ‘by=sample’ parameter as input. |
| **1.4** Could you please clarify the precise formula employed for determining rounding errors and error recovery in the Method section? I suppose each gene in each cell should have a discrepancy. Did you sum them up or calculate the average value of them? | Sorry for the confusion! The error metrics we use do not sum up or calculate the average values. So an error is measured for a single value for one gene in one cell. We are now providing formulas for calculating rounding error and recovery error in the Methods section, see P12L464.<br><br>When benchmarking denormalisation for scRNA-seq data, two measures can be used: rounding error and recovery error. Rounding error measures the discrepancy between the denormalised values and their rounded counterparts. After denormalisation, the expected outcome is that the denormalised values approximate integers. Rounding error quantifies the extent to which the denormalised values deviate from integers. To calculate rounding error, the difference between each denormalised value and its rounded value is computed, see equation (9) below. Recovery error evaluates the difference before denormalisation and after re-normalising the denormalised values (values after scDenorm, **Fig. 3a**). To calculate recovery error, the difference between each normalised value and its re-normalised value is computed, see equation (10) below.<br><br>Specifically, we assume x is the normalised value (a single value for one gene in one cell), y is the denormalised value after scDenorm, and z is the re-normalised value from denormalised value (y). The rounding error is calculated as the difference between the denormalised value (y) and its rounded value, equation (9):<br><br>$$rounding\_error = round(y) - y \quad (9)$$ |

| | |
|---|---|
| | The recovery error is calculated as the difference between the normalised value (x) and the re-normalised value (z), equation (10):<br><br>$$recovery\_error = x - z \quad (10)$$<br><br>In certain cases, not all cells can be successfully denormalised due to poor sequencing quality, or a low number of genes. To evaluate denormalisation in such situations, we define success rate as the percentage of successfully denormalised cells, equation (11).<br><br>$$success\ rate = N_{success}/N_{total} \quad (11)$$<br><br>Nsuccess is the number of successfully denormalised cells, while Ntotal is the total number of cells. |
| **1.5** In Figure 1c, the author said they perform integration using Harmony, scanorama and BBKNN. The results for Harmony are in Figure 1c. The results for BBKNN are in Supplementary Fig 1. Where is the results for scanorama? In addition, for enhanced clarity, it would be beneficial to include the name of the integration algorithm not only in the figure legend but also directly within the figure itself. | Sorry for the misunderstanding. We are now including the scanorama result in **Supplementary Fig. 1c**. The result does not show clear cell type distribution as expected, indicating the necessity of denormalisation.<br><br>Following this reviewer's suggestion, we have now updated the figure labels and figure legends to be self-explainable. See **Supplementary Fig. 1**. |

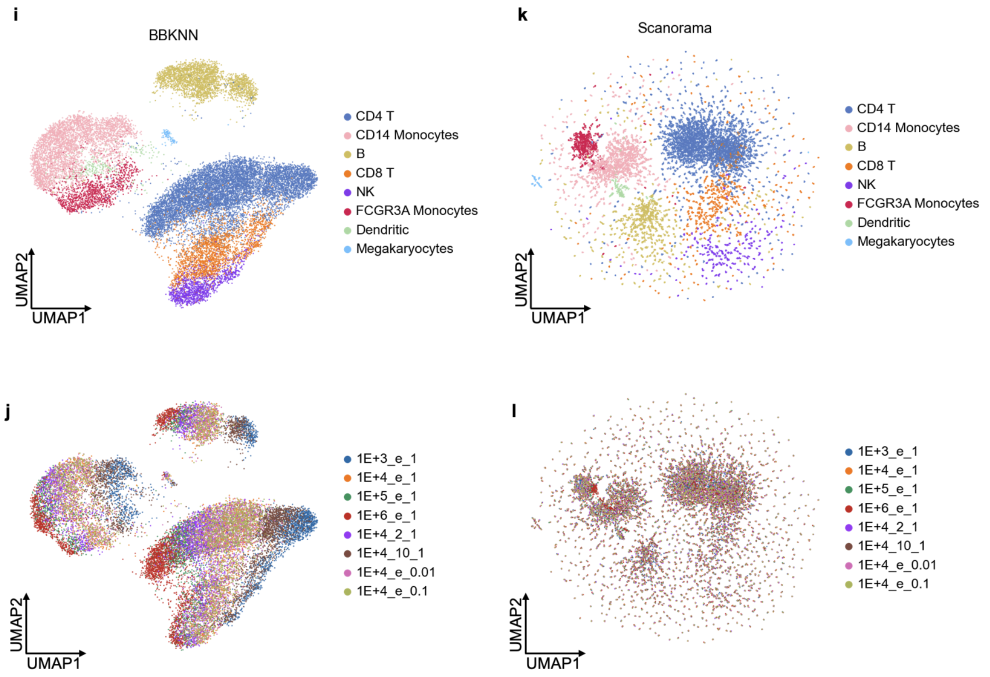


**Supplementary Fig. 1 Inconsistent data normalisation generates bias in data integration**

**i-j**, UMAP plots show the BBKNN-integrated result on the data obtained from different normalization methods, coloured by different cell types (**i**) and different normalization methods (**j**).

**k-l**, UMAP plots show the scanorama-integrated result on the data obtained from different normalization methods after Scanorama integration, coloured by different cell types (**k**) and different normalization methods (**l**).

| | |
|---|---|
| **1.6** In the figure legends for 1d, it appears that 1d differs significantly from 1b. Therefore, did you mean the UMAP plots are the same as (c)? | Sorry for the mistake. Yes, the UMAP plots in 1d should be the same as 1c, not 1b. We have now corrected the figure legend. |
| **1.7** Figure 2 presents the method and formula, which are already described in detail in the Method section. I think it is redundant to include those formulas in a main figure. Therefore, I propose placing them in the supplementary material and replacing Figure 2 with a figure showcasing results. | Thanks! Following the suggestion, we have now moved **Fig. 2** to **Supplementary Fig. 3**. |
| **1.8** In Figure 3a,d,g, I'm wondering how to select the three cells for demonstration? Do the three cells have something special? | Thanks. The three cells shown in Figures 3a, 3d, and 3g are the first three cells from the dataset. As all the cells follow the same distribution, we just randomly selected them for illustration. We have now clarified this point at P4L148:<br><br>Should any of the cells in the dataset follow the same distribution.<br><br>Here, we randomly selected other three cells in the dataset and show the same distribution as in **Revision Fig. 2**. |

| | |
|---|---|
| | 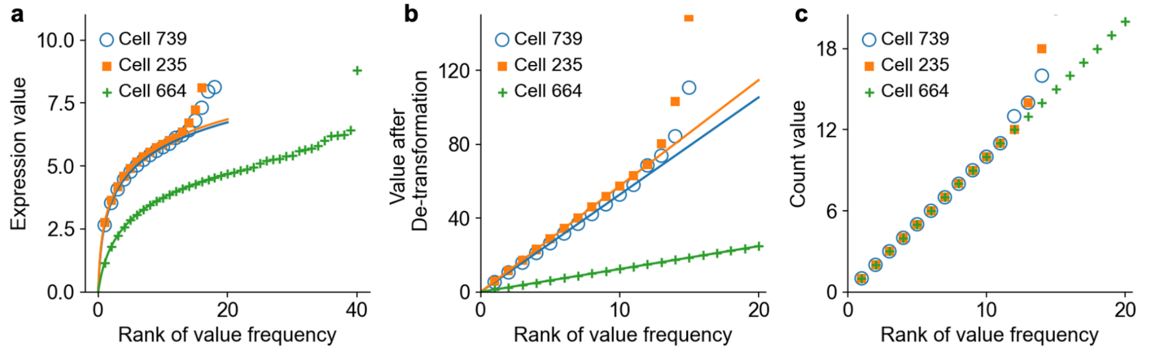 <br>**Revision Fig, 2 the relationship between the value of gene expression and the rank of their frequencies during denormalisation process**<br>**a,** The scatter plot of the rank and its value of gene expression in 3 random cells. Each colour for a cell.<br>**b,** The scatter plot of the rank and its value of gene expression in the above cells after de-transformation.<br>**c,** The scatter plot of the rank and its value of count in the above cells after denormalisation. |
| **1.9** In Figure 4b, it is unclear what the color represents. In addition, the number of cells in each dataset might also be useful information. Could you use another y axis to show the number of single cells for each dataset. | Thank! In Figure 4b, the different colors represent the different normalization parameters that were detected using scDenorm. We have now updated the figure legends for clarity, see P21L740:<br><br>Following this reviewer's suggestion, we have updated the new **Fig. 3b (previous Fig. 4b)** to show the number of cells for each dataset on an additional y-axis. |

<table>
<tr>
<td></td>
<td>
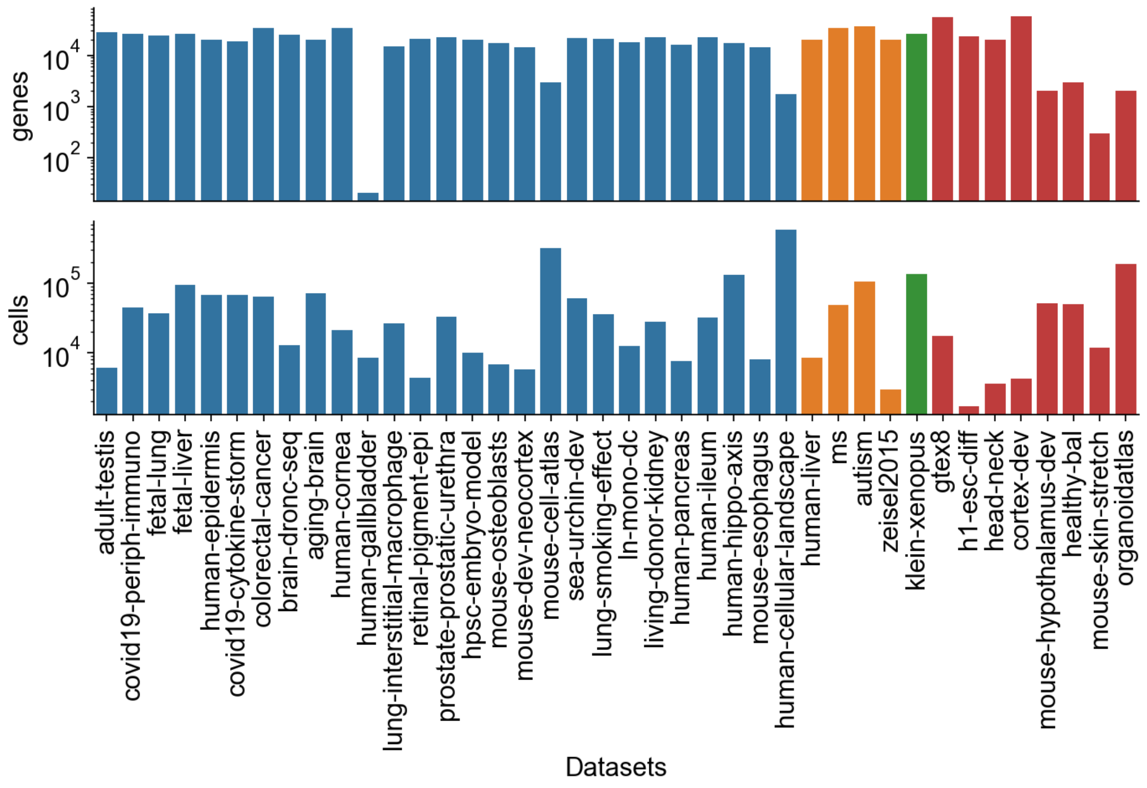


**Fig. 3b,** Two barplots show the number of genes (top) and number of cells (bottom) for the collected UCSC datasets. The x-axis shows the datasets by name, while the y-axis shows the log-scaled number of genes (top) and the log-scaled number of cells (bottom). The colours represent the different parameters of the delta normalisation. Blue and orange are natural base (e) and base=2 respectively, green represents data without log-transformation. Red shows non-delta method normalisation cases, which could not be denormalised by scDenorm.
</td>
</tr>
<tr>
<td>**1.10** The figure 4d seems unclear.</td>
<td>Sorry for the confusion. We have separated the figure 4d into different panels (updated</td>
</tr>
</table>

What is the relationship of de-normalised values in figure legend and total value in figure 4d (left)? The author wrote "The scatter plot of the relationship between the de-normalised values and rounding errors (left)" in the legend, but I can not find rounding errors in the left panel. In addition, the author wrote "The distribution of scaling factor (left) and rounding errors (right) for each of the de-normalised datasets (middle)." in the legend, but the right panel refer to recovery errors.

Fig. 3d, 3e, 3f) and updated the figure legend for clarity. Please refer to the revised version in P21L749:

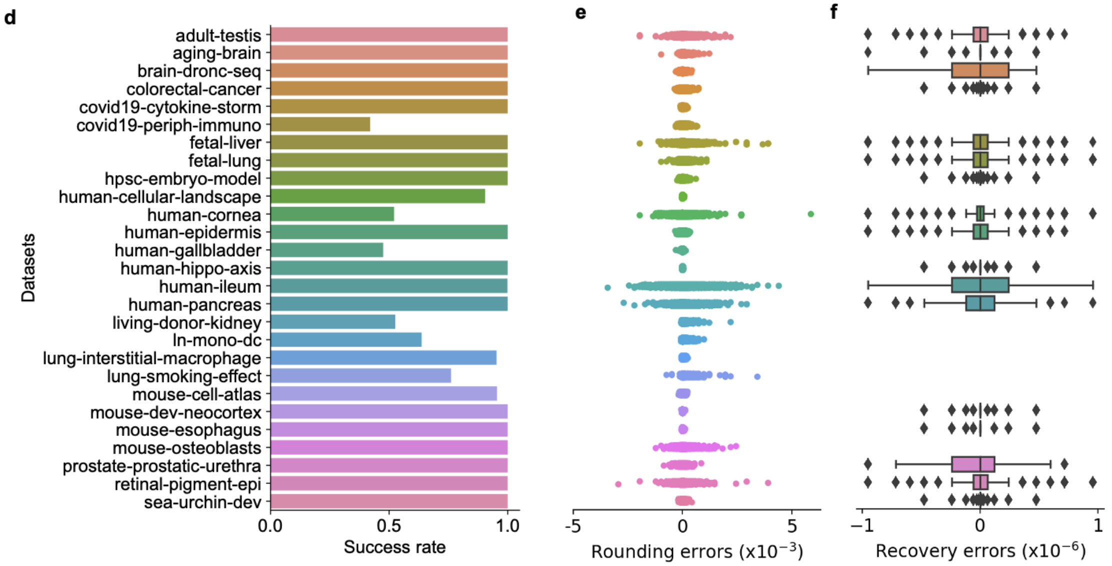


**Fig. 3 Performance evaluation of scDenorm on normalised scRNA-seq data from UCSC Cell Browser**

**d,** The bar plot shows the distribution of the success rate (see **Methods**) across the 27 datasets that were normalised with natural logarithmic transformation.

**e,** The jitter plot shows the distribution of rounding errors observed in the denormalised datasets. The x-axis is the rounding error, while the y-axis shows the same datasets as panel (**d**).

**f,** The box plot shows the distribution of recovery errors after re-normalization with the parameters of target sum, pseudo count and logarithm base as 1e4, 1 and natural base (e), respectively. The x-axis is the recovery error, while the y-axis shows the datasets in the same order as in panel (**d**).

<table>
<tr><td>1.11 In Figure 5f, I'm wondering how to perform UMAP for both Origin and HVG since they have different number of genes. Did you perform UMAP on Origin or HVG separately and then plot together? In addition, I suppose the number of cells in Origin, HVG and Floating should be equal, but it seems that the number of green dots is much greater than blue and orange dots. Or the colors you selected influence the demonstration?</td><td>Sorry for the confusion. We performed the UMAP on the intersection of genes between Origin and HVG. The number of cells in Origin, HVG, and Floating are the same. Due to the layered visualization of different color dots in the UMAP, it can give the impression that the green dots (HVG) are more than the blue (Origin) and orange (Floating) dots.<br><br>To clarify, we separated the above UMAP and showed the UMAPs by Origin, HVG, and Floating in Revision Fig. 3. As we can see that the three data representations show similar UMAP distributions.<br><br>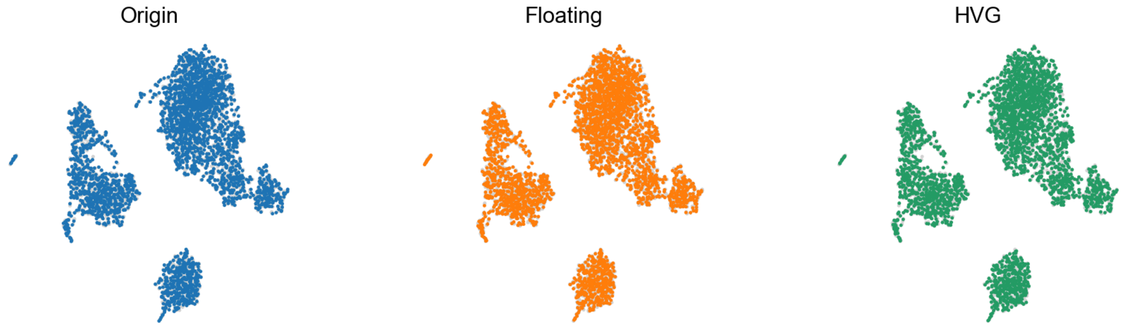<br><br><br>Revision Fig, 3 UMAP plots with different colours to represent different cases, including original data (blue), denormalised data after converting to floating 16 (yellow), and denormalised data after selecting 2,000 highly variable genes (green).</td></tr>
<tr><td>Reviewer 2</td><td></td></tr>
<tr><td>2.0 In the current submission, Huang et al. introduced "scDenorm: A Denormalization Tool for Integrating Single-cell Transcriptomics Data." While the reviewer appreciates the effort and time invested in your research, after careful evaluation, they</td><td>Thanks! Following the reviewers' suggestions, we have now made significant revisions of our manuscript, including more complete benchmarking, improved methodology clarity and systematic text/figure revision. Please see answer to comment 1.0 for details.</td></tr>
</table>

| | |
|---|---|
| feel that the current submission requires significant revision to meet GB standards. | |
| **2.1** First, the novelty of the study appears limited. While the concept of reversing normalized single-cell omics data to raw counts is innovative, the manuscript does not sufficiently demonstrate how this approach offers a significant advancement over existing methods. Highlighting unique aspects or potential breakthroughs facilitated by scDenorm would strengthen the manuscript's impact. | Sorry for the confusion. This comment conflicts with reviewer 1's **comment 1.0**, as "The idea of denormalisation is innovative and this algorithm is relatively concise and should be quite effective. " To clarify the novelty of our work and avoid potential confusion, we would like to highlight:<br><br>1. To the best of our knowledge and literature search, there are no other existing methods, which can denormalise single-cell data to raw counts.<br><br>We have also clarified it in the manuscript, P2L75.<br><br>Yet, there is no tool available to meet this urgent need.<br><br>2. For the unique aspects of scDenorm, a) denormalising data may avoid false discoveries in downstream analysis, see answer to **comment 1.1**; b) it can process large-scale dataset, facilitating the integration of atlas scale datasets, see P5L171.<br><br>To evaluate the performance of scDenorm in real scenarios, 40 processed datasets (**Supplementary Table 1**) from the UCSC Cell Browser[24] were used as test data covering a good variety of species, tissues and sequencing techniques (**Fig. 3b**). Denormalisation performance was evaluated by two metrics: 1) rounding error, defined as the difference between a value in the denormalised matrix and its nearest integer (round value), and 2) recovery error, defined as the difference between a value in the normalised matrix and its corresponding value in the denormalised matrix after renormalisation (**Fig. 3a**, See **Methods**). 32 out of the 40 test sets were successfully denormalised (**Fig. 3c**, **Supplementary Table 1**), while the 8 unsuccessful cases were normalised as TPM (Transcripts per Million), log2(FRPM) or by scTransform[32] method rather than the delta method (**Supplementary Table 1**). The mean versus variance |

| | |
|---|---|
| | distribution confirms a negative binomial distribution after denormalisation, indicating the successful denormalisation (**Supplementary Fig. 5**). The rounding errors, which are positively correlated to the expression value (**Supplementary Fig. 6**), are consistently below 0.005 (**Fig. 3e**). For recovery error, the absolute values are below $10^{-6}$ in 27 datasets normalised with natural logarithmic transformation (**Fig. 3d, Supplementary Table 1**), indicating a good accuracy of scDenorm (**Fig. 3f**). Further benchmarking of the denormalisation on 60 datasets (**Supplementary Table 2**) from the Brain Cell Atlas[30] show similar results (**Supplementary Fig. 7**). In addition, scDenorm shows a linear computational time complexity and memory usage with increasing number of cells and genes, demonstrating a high computational efficiency and scalability (**Supplementary Fig. 4c,d**). |

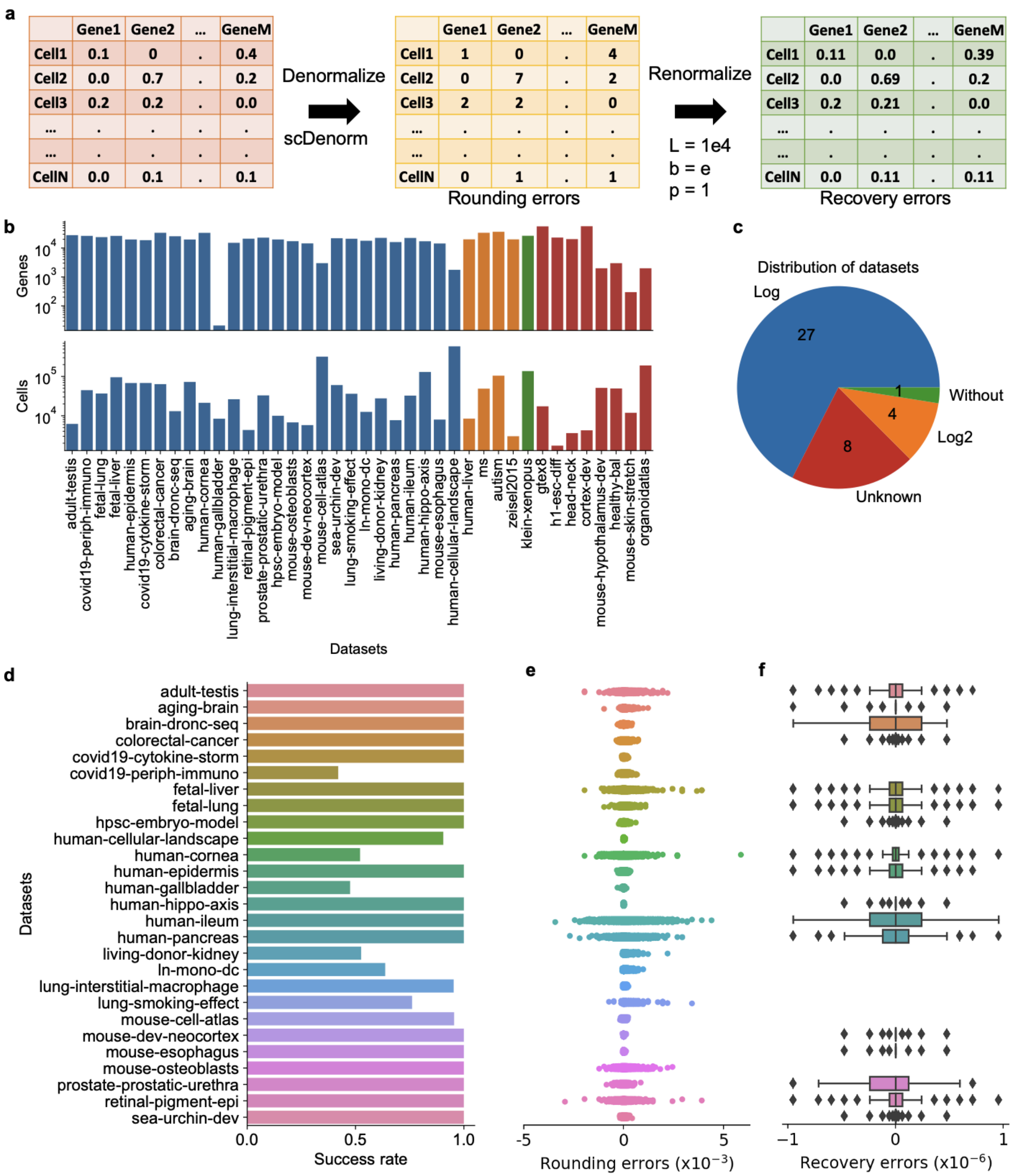
a
Gene1
Gene2
...
GeneM
Cell1
Cell2
Cell3
CellN
Denormalize
scDenorm
Rounding errors
Renormalize
L = 1e4
b = e
p = 1
Recovery errors
b
Genes
Cells
Datasets
adult-testis
covid19-periph-immuno
fetal-lung
fetal-liver
human-epidermis
covid19-cytokine-storm
colorectal-cancer
brain-dronc-seq
aging-brain
human-cornea
human-gallbladder
lung-interstitial-macrophage
retinal-pigment-epi
prostate-prostatic-urethra
hpsc-embryo-model
mouse-osteoblasts
mouse-dev-neocortex
mouse-cell-atlas
sea-urchin-dev
lung-smoking-effect
ln-mono-dc
living-donor-kidney
human-pancreas
human-ileum
human-hippo-axis
mouse-esophagus
human-cellular-landscape
human-liver
ms
autism
zeisel2015
klein-xenopus
gtex8
h1-esc-diff
head-neck
cortex-dev
mouse-hypothalamus-dev
healthy-bal
mouse-skin-stretch
organoidatlas
c
Distribution of datasets
Log
27
Without
1
4
Log2
8
Unknown
d
Datasets
Success rate
e
Rounding errors (x10$^{-3}$)
f
Recovery errors (x10$^{-6}$)

**Fig. 3 Performance evaluation of scDenorm on normalised scRNA-seq data from UCSC Cell Browser**

**a,** The diagram illustrates the workflow of scDenorm to evaluate denormalisation using the rounding and recovery error. The processed matrix (left) deposited in the UCSC database is first denormalised (middle) with scDenorm to calculate the rounding error (see **Methods**). Subsequently, the denormalised matrix (middle) is re-normalised (right) to measure the recovery error (see **Methods**). The values of target sum and pseudo count normalisation parameters are 1e4 and 1, respectively.

**b,** Two barplots show the number of genes (top) and number of cells (bottom) for the collected UCSC datasets. The x-axis shows the datasets by name, while the y-axis shows the log-scaled number of genes (top) and the log-scaled number of cells (bottom). The colours represent the different parameters of the delta normalisation. Blue and orange are natural base (e) and base=2 respectively, green represents data without log-transformation. Red shows non-delta method normalisation cases, which could not be denormalised by scDenorm.

**c,** The pie chart shows the distribution of the number of datasets classified by different base values, detected by scDenorm. The colours are the same as shown in panel (**b**). Unknown represents non-delta method normalisation cases.

**d,** The bar plot shows the distribution of the success rate (see **Methods**) across the 27 datasets that were normalised with natural logarithmic transformation.

**e,** The jitter plot shows the distribution of rounding errors observed in the denormalised datasets. The x-axis is the rounding error, while the y-axis shows the same datasets as panel (**d**).

**f,** The box plot shows the distribution of recovery errors after re-normalization with the parameters of target sum, pseudo count and logarithm base as 1e4, 1 and natural base (e), respectively. The x-axis is the recovery error, while the y-axis shows the datasets in the same order as in panel (**d**).

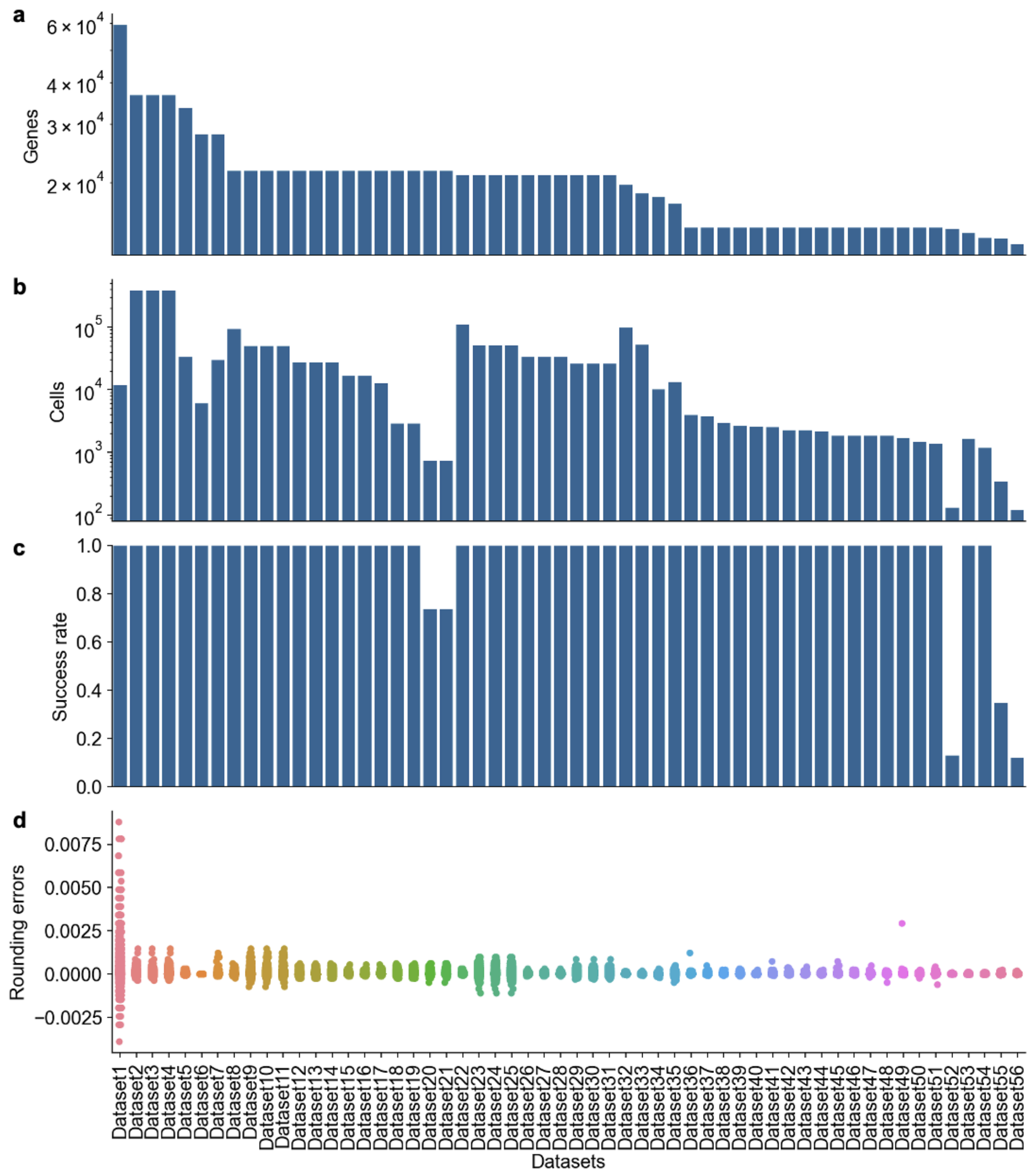

a
Genes
6 × 10^4
4 × 10^4
3 × 10^4
2 × 10^4
b
Cells
10^5
10^4
10^3
10^2
c
Success rate
1.0
0.8
0.6
0.4
0.2
0.0
d
Rounding errors
0.0075
0.0050
0.0025
0.0000
−0.0025
Dataset1 Dataset2 Dataset3 Dataset4 Dataset5 Dataset6 Dataset7 Dataset8 Dataset9 Dataset10 Dataset11 Dataset12 Dataset13 Dataset14 Dataset15 Dataset16 Dataset17 Dataset18 Dataset19 Dataset20 Dataset21 Dataset22 Dataset23 Dataset24 Dataset25 Dataset26 Dataset27 Dataset28 Dataset29 Dataset30 Dataset31 Dataset32 Dataset33 Dataset34 Dataset35 Dataset36 Dataset37 Dataset38 Dataset39 Dataset40 Dataset41 Dataset42 Dataset43 Dataset44 Dataset45 Dataset46 Dataset47 Dataset48 Dataset49 Dataset50 Dataset51 Dataset52 Dataset53 Dataset54 Dataset55 Dataset56
Datasets

| | |
|---|---|
| | **Supplementary Fig. 7 Benchmark analysis on the datasets from Brain Cell Atlas**<br>**a,** The bar plot shows the number of genes in each dataset. The x-axis is the name of datasets (same as **d**), and the y-axis is the log-scaled number of genes.<br>**b,** The bar plot shows the number of cells in each dataset. The x-axis is the name of datasets (same as **d**), and the y-axis is the log-scaled number of cells.<br>**c,** The bar plot shows the success rate for each dataset. The x-axis is the name of datasets (same as **d**), and the y-axis is the success rate (see **Method**).<br>**d,** The jitter plot shows the distribution of rounding errors observed in the denormalised datasets from the Brain Cell Atlas. The x-axis is the name of datasets, and the y-axis is the rounding error. |
| **2.2** Second, the reviewers believe that more extensive validation is necessary to establish the biological relevance of scDenorm. This includes providing detailed case studies or examples where scDenorm effectively mitigates biases in downstream analyses and contributes to new biological insights. Without such validation, it is challenging to assess the practical utility and reliability of the proposed method. | Thanks for the suggestion! Please also see the answer to **comment 1.1**.<br><br>Besides the large-scale benchmarking test with the UCSC database, we have now added further large-scale benchmark test on 65 human brain datasets (see **Supplementary Fig. 7**) and included three real-world study cases (COVID-19 PBMC dataset (see **Fig. 5**), human skin dataset (see **Fig. 6**), and human prefrontal cortex dataset (see **Supplementary Fig. 11**) to demonstrate the biological relevance of scDenorm in mitigating biases in downstream analyses.<br><br>Specifically, scDenorm effectively reduced biases in cell type annotation and differential expression analysis, leading to more accurate results in enrichment analyses, see **Fig. 5 and Fig. 6**. This not only enhances the reliability of our findings but also contributes to new biological insights, particularly by alleviating unnecessary biases in atlas-level data integration. |
| **2.3** Third, the manuscript requires more extensive benchmarking against a broader range of normalization methods and datasets. A thorough comparison with current | Sorry for the confusion. We would like to clarify that there are no other existing methods of the same function available, see answer to **comment 2.1**.<br><br>Following this reviewer's suggestion, we are now including more extensive benchmarking of the method. |

| | |
|---|---|
| state-of-the-art techniques is essential to convincingly demonstrate the robustness and accuracy of scDenorm. This benchmarking should include performance metrics such as computational efficiency, scalability, and the ability to preserve biological signal integrity. | 1. For computational efficiency and scalability, our result indicates that scDenorm demonstrates high computational efficiency and scalability, with a linear relationship between computation time complexity and memory usage with the number of cells and genes (see **Supplementary Fig. 4c,d**).<br><br>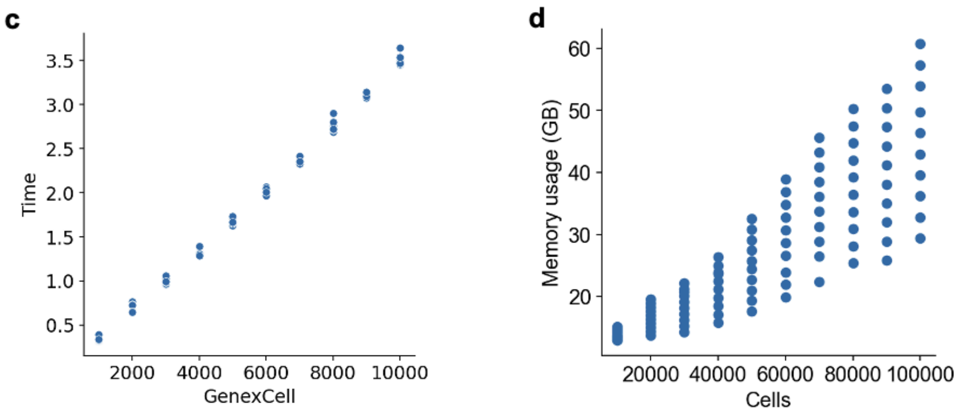<br><br><br>**Supplementary Fig. 4 The computational efficiency of scDenorm**<br>**c**, The distribution of execution times for denormalisation (y-axis, the unit is second) across cells with genes ranging from 2000 to 10000 (x-axis).<br>**d**, The distribution of memory usage during denormalisation (y-axis, the unit is GB) across cells ranging from 10000 to 100,000 (x-axis).<br><br>2. For the ability to preserve biological signal integrity, we have now included three real-world study cases (COVID-19 PBMC dataset (see **Fig. 5**), human skin dataset (see **Fig. 6**), and human prefrontal cortex dataset (see **Supplementary Fig. 11**) to demonstrate the biological relevance of scDenorm in mitigating biases in downstream analyses. See answer to **comment 2.2**. |

| | |
|---|---|
| | 3. For the robustness and application scope, we have benchmarked scDenorm using 40 datasets from the UCSC Cell Browser and 60 datasets from the Brain Cell Atlas. Over 88% of these datasets were normalised by delta method, while scDenorm effectively recovered raw counts for all of these delta method normalised datasets (see **Fig. 4, Supplementary Fig. 7**). Therefore, we expect scDenorm applies to a wide range of datasets. |
| **2.4** Fourth, technical clarity is a concern. Sections detailing the algorithm's implementation and theoretical foundations need to be more comprehensible. Providing clearer explanations and potentially including visual aids or supplementary materials could help in making the methodology more accessible to a wider audience. | Thanks. We have now systematically revised the Methods sections (including the algorithm's implementation, evaluation metrics and dataset processing) for clarification. see P9L330:<br><br>Additionally, we have included a schematic illustration of the algorithm (see **Supplementary Fig. 3**) and a visual representation of the data changes during the denormalisation process (see **Fig. 2**) to better illustrate the idea and mechanism of denormalisation. Please find **Supplementary Fig. 3** and **Fig. 2** as below: |

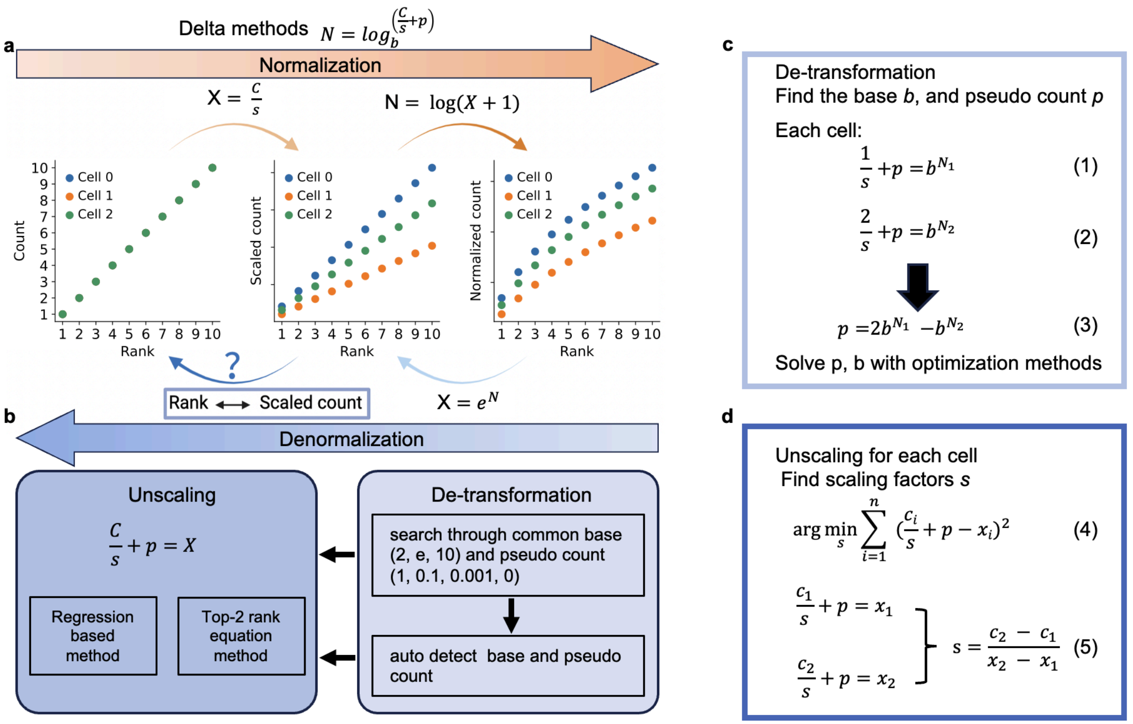


**Supplementary Fig. 3 Schematic representation of scDenorm algorithm**

a, Standard pre-processing: routinely droplet-based single-cell data are first normalised by a size factor (s, usually total count with a library size factor) with and pseudo count (p) secondly transformed by a logarithmic function with a base (b) for downstream analysis.

b, Denormalisation: scDenorm performs the reverse of the task above. First it reverses the logarithmic transformation by searching for common bases (e.g., 2, e, 10) and common pseudo counts (e.g., 0.01, 0.1, 1), or auto detect base and pseudo count. Then, it reverses the scaling by determining the scaling factor for each cell.

c, The formulas and derivations of automatic detection method for base and pseudo count during the de-transformation process. b is the base, p is the pseudo count, s is the scaling factor, and N1 and N2 are the most and second frequency values for each cell.

| | |
|---|---|
| | d, The formulas of solving the scaling factor for each cell during the unscaling process. Equation (4) is the formula of the regression based method, and equation (5) is the formula of the top-2 rank equation method. ci are the ranks of the frequency of normalised values ( c1 is 1, and c2 is 2, etc). xi are the values sorted by frequency in a decreasing order after de-transformation. |

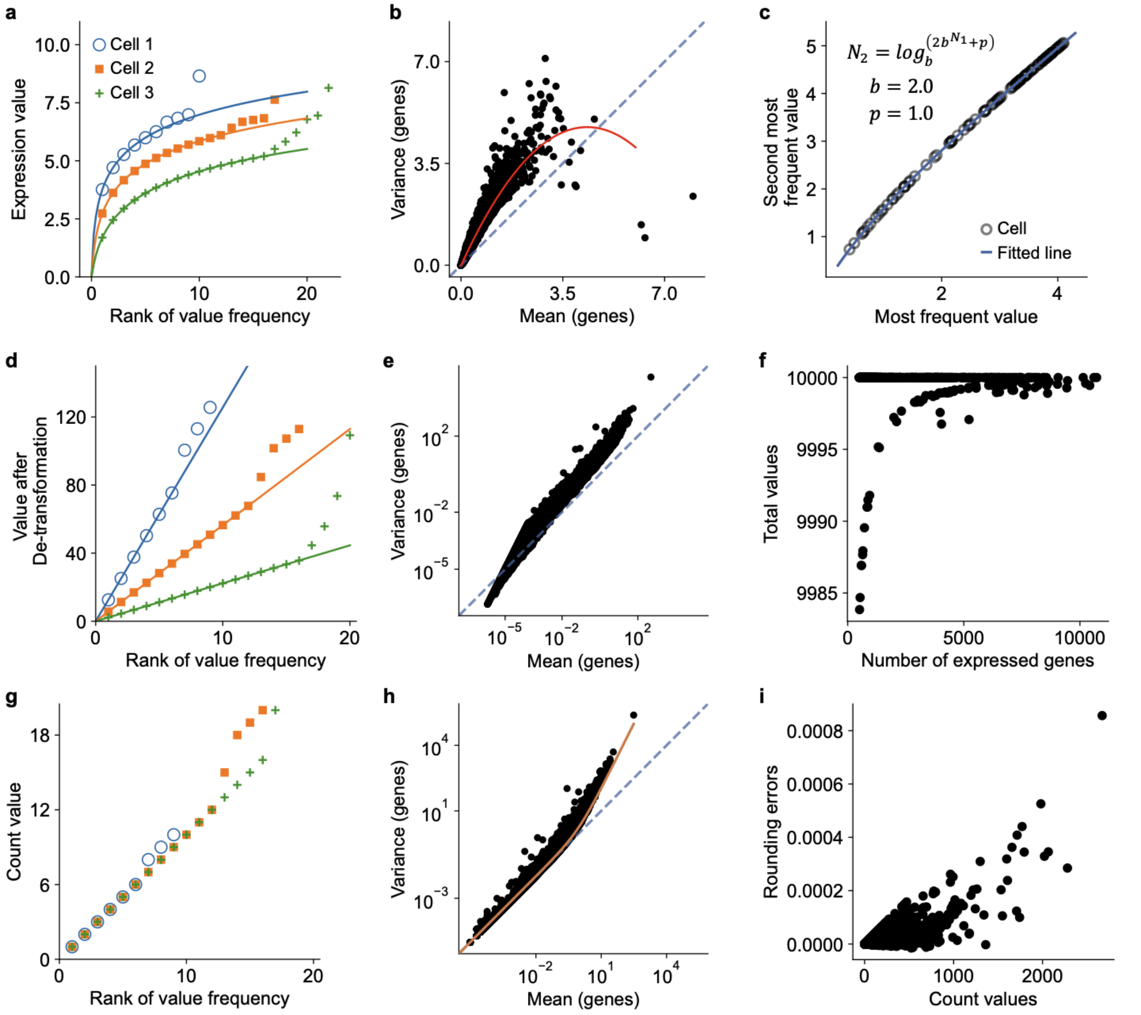


**Fig. 2 Evaluation of scDenorm on normalised scRNA-seq data with known raw counts**

| | |
|---|---|
| | **a,** The scatter plot shows the distribution between expression values and their ranks of frequencies in three example cells. Each dot is an expression value and its rank of frequency in the cell, different cells are shown in different shape and colour.<br>**b,** the scatter plot shows the distribution between the log-transformed mean expression (x-axis) and the log-transformed variance (y-axis) for each gene in the gene expression matrix from Velmeshev et al dataset. The diagonal line (x=y) is shown in blue.<br>**c,** The scatter plot shows the distribution between the most and second most frequent values in different cells, displaying each cell as a dot. The blue curve shows the fitted equation derived from the equation (4) in **Methods**, with base value (b) equal to 2 and pseudo count (p) equal to 1. $N_1$ and $N_2$ are the most and second most frequent values in cells.<br>**d,** The scatter plot shows the distribution between expression values and their ranks of frequencies in the three example cells after de-transformation, coloured in the same manner as panel (**a**).<br>**e,** The scatter plot shows the relationship between the mean expression (x-axis) and variance (y-axis) for each gene after de-transformation.<br>**f,** The scatter plot shows the distribution between the number of genes and the target sum (sum of all expression values) in the cell after de-transformation.<br>**g,** The scatter plot shows the distribution between expression values and their ranks of frequencies in the three example cells after unscaling, displaying the 'count-rank' distribution.<br>**h,** The dot plot shows the distribution between the mean expression (x-axis) with the variance (y-axis) for each gene in the count matrix.<br>**i,** The scatter plot shows the distribution between the count values and the rounding errors after denormalisation. Each dot represents a count value in a cell and its rounding error. |
| **2.5** Finally, the language and overall presentation of the manuscript need improvement. This includes refining the writing for clarity, coherence, and | Thanks. We have tried our best to improve the language and overall presentation. We have checked the language with both Grammarly (grammarly.com) and deepl (deepl.com). |

| | |
|---|---|
| conciseness, as well as ensuring proper use of technical terminology. Enhancing the manuscript's readability will facilitate better understanding and appreciation of your work by the broader scientific community. | Please let us know if further improvement is required. We are committed to improving the clarity, coherence, and conciseness of our writing to enhance its readability for the broader scientific community. |
